# Dorsal-to-ventral neocortical expansion is physically primed by ventral streaming of early embryonic preplate neurons

**DOI:** 10.1101/601617

**Authors:** Kanako Saito, Mayumi Okamoto, Yuto Watanabe, Namiko Noguchi, Arata Nagasaka, Yuta Nishina, Tomoyasu Shinoda, Akira Sakakibara, Takaki Miyata

## Abstract

Mammalian neocortex exhibits a disproportionally “luxurious” representation of somatotopies in its lateral region, which depends on dorsal-to-ventral expansion of the pallium during development. Despite recent studies elucidating the molecular mechanisms underlying the cortical arealization/patterning, we know very little about how the cortex expands ventrally and the nature of the underlying force-generating events. We found that neurons born earliest (at embryonic day 10 [E10]) in the mouse pallium migrated ventrally and then extended corticofugal axons, which together formed a morphogenetic flow of the preplate that persists until E13. These neurons exerted pulling and pushing forces at the process and the soma, respectively. Ablation of these E10-born neurons attenuated both deflection of radial glial fibers (by E13) and extension of the cortical plate (by E14), which should occur ventrally, and subsequently shrank the postnatal neocortical map dorsally. This previously unrecognized preplate stream physically primes neocortical expansion and somatotopic map formation.

## Introduction

The lateral part of the adult mammalian neocortex, including the parietal lobe where the homunculus or mouseunculus is mapped, is superficial to the striatum (Figure 1A). Embryonically, the pallium (neocortical primordium) is dorsal to the ganglionic eminences (striatal primordia), with the palliostriatal (PS) angle serving as a morphological demarcation along the ventricular surface. By contrast, the adult lateral neocortex is ventral to the PS angle, indicating that the pallium undergoes ventral expansion/growth in order to “ride on” the striatum. Clonal analyses revealed a greater tangential dispersion of neurons in the lateral part than in the medial part (Tan et al., 1995; Gao et al., 2014). One prominent explanation for this ventral neocortical expansion is that radial glial fibers, which are used for neuronal guidance (Rakic, 1972), exhibit an extensively curving and divergent alignment pattern in the lateral part (Misson et al., 1988; 1991) (Figure 1B), which can be coupled with tangential translocations of neurons from one radial fiber (RF) to another (O’Rourke et al., 1992; Tabata and Nakajima, 2003). This ventral curving/deflection of RFs seems to be specific to mammals, and is absent in the chick (Striedter and Beydler, 1997). In addition, the fact that neuron production by progenitors begins and proceeds earlier in the lateral/ventral part than in the dorsomedial parts (Hicks and D’Amato, 1968; Smart and Smart, 1977) allows us to imagine that tissue volume can easily increase laterally or ventrally. There remains, however, an additional model that has not been previously tested due to technical difficulties: dorsal-to-ventral migration of neurons born very early in the pallium could contribute to the ventral-dominant neocortical expansion, either directly or by causing (rather than resulting from) the curvature of RFs.

**Figure 1.**
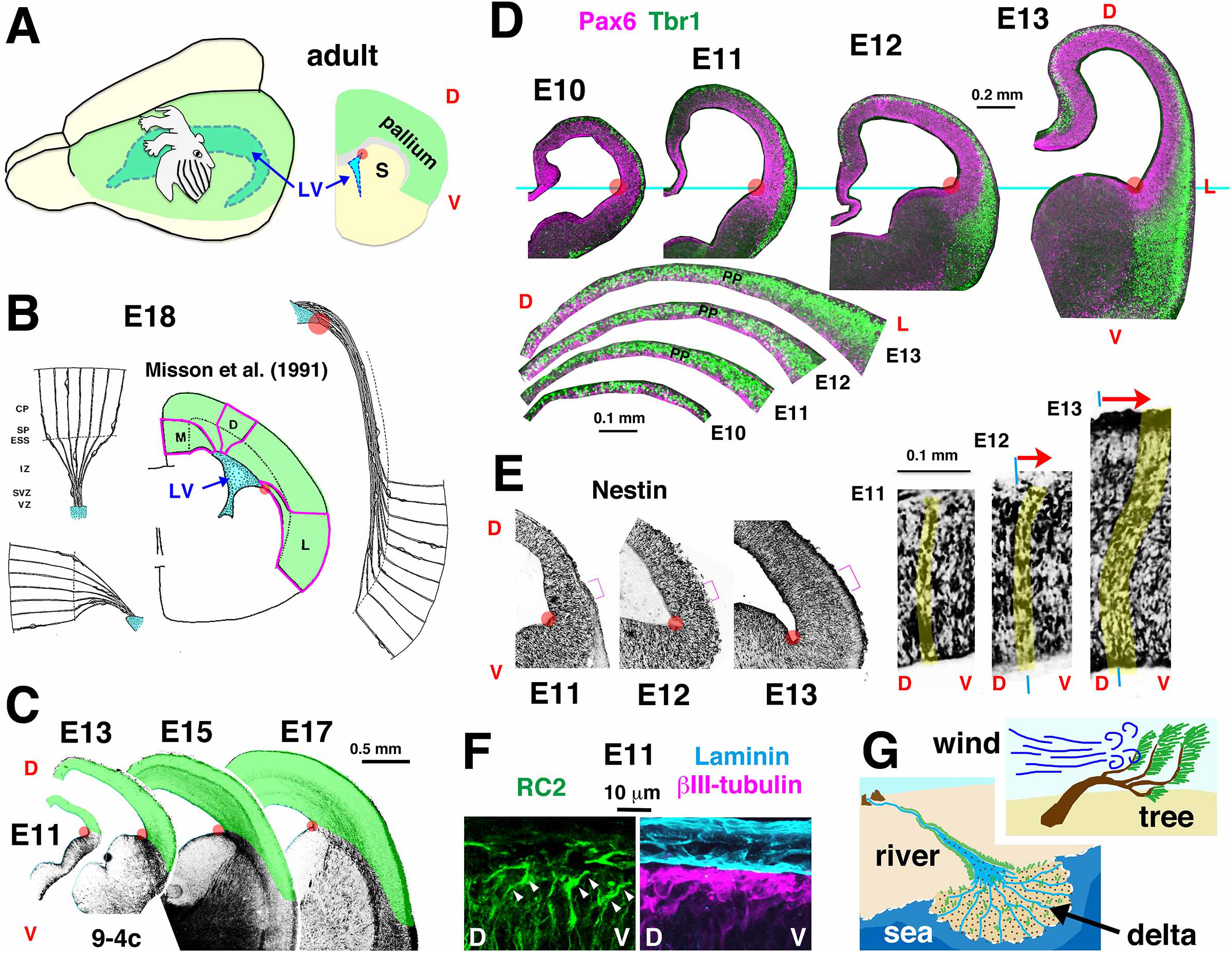
Neocortical expansion and deflection of RFs, which occur ventrally during the embryonic period. (A) Adult mouse neocortex (green), with a somatosensory mouseunculus, spread ventral to the pallio-striatal angle (PS angle, red spot). LV, lateral ventricle. S, striatum. D, dorsal. V, ventral. (B) Asymmetric ventralward deflection of RFs in the lateral (L) region of E18 neocortex (from Misson et al., 1991). M, medical. CP, cortical plate. SP, subplate. ESS, external sagittal stratum. IZ, intermediate zone. VZ, ventricular zone. (C) 9-4c immunoreactivity in the pallio-subpallial boundary (PSP boundary). (D) Superimposed images of anti-Pax6 and anti-Tbr1 immunoreactivity, obtained from two adjacent sections. L, lateral. PP, preplate. (E) Anti-Nestin staining. (F) RC2, anti-Laminin, and anti–βIII-tubulin immunostaining. Distal RFs were ventrally deviated (arrowhead). (G) Physical analogies to facilitate understanding of possible mechanisms underlying neocortical expansion and RF deflection, which occur ventrally during the early embryonic period.

Dorsal-to-ventral tangential migration into the pallium has been observed in Cajal–Retzius (CR) cells derived from the hem region (Takiguchi-Hayashi et al., 2004; Borrell and Marin, 2006; Yoshida et al., 2006). The early emergence of CR cells underlies their histogenetic influence on later-born neurons (Ogawa et al., 1995; de Frutos et al., 2016), which is assisted by their tangential dispersion (Bielle et al., 2005; Barber et al., 2015). By contrast, the majority of subplate (SP) neurons, another developmentally influential type of early-born neurons (McConnell et al., 1989; Ghosh et al., 1990; Ohtaka-Maruyama et al., 2018; reviewed in Allendoerfer and Shatz, 1994; Hoerder-Suabedissen and Molnár, 2015), are thought not to migrate tangentially from their origins in the dorsal pallium (reviewed in Barber and Pierani, 2016). However, recent studies have shown that SP neurons come from an extracortical (rostromedial) origin via a medial-to-lateral influx (Pedraza et al., 2014; García-Moreno et al., 2018), and that a fraction of neurons born in the ventricular zone (VZ) of the pallium and PS interface migrate ventrally to contribute to the olfactory cortex, amygdala, striatum, or other forebrain structures (Tomioka et al., 2000; Hamasaki et al., 2001; Hirata et al., 2002; García-Moreno et al., 2008; Cocas et al., 2009; Pombero et al., 2011). Together, these observations prompted us to carefully investigate whether pallium-derived SP neurons also migrate ventrally.

In this study, we performed site-directed labeling of the mouse dorsal pallium via *in utero* electroporation (IUE) at E10, followed by tracking both *in vivo* and in slice culture. We found that the earliest pallial-born neurons that constitute the preplate (PP) (Rickmann et al., 1977; Valverde et al., 1995) (also known as the primordial plexiform layer [Marin-Padilla, 1971]), which abundantly contained SP-destined neurons, underwent a massive ventral migration during the E11–12 period that is dependent on Rac1 and N-Cadherin. These cells exhibited a locomotion-like migratory mode and exerted traction (at the leading process) as well as pushing (at the soma). This PP stream continued (albeit less rapidly) until E13, when multiple somal behaviors reflecting heterogeneous neuronal morphologies (including nucleokinesis into corticofugal axons), rather than the preexisting locomotion mode, drove a collective, tissue-level flow into which E11- or E12-born neurons entered in an oblique (i.e., ventrally deviated) manner in order to initiate the cortical plate (CP). E10 IUE-mediated reduction of these PP neurons attenuated the ventral deflection of RFs at E13 and the ventral extension of CP at E14, resulting in a dorsal shift of the neocortical map, including somatosensory barrels, in the early postnatal period. Thus, this ventral PP stream from E11 to E13, a long-sought ‘missing piece’ in our understanding of neocortical ontogeny, plays a key mechanical role in the expansion of the mammalian neocortex, revealing the unexpected reliance of the RF-based protomap mechanism (Rakic, 1988) on early tangential (frontier-advancing) neuronal migration.

## Results

### Ventral expansion of the neocortical region begins in the early embryo with ventral deflection of the distal RFs

To morphometrically analyze the lateral/ventral expansion of the neocortical region during development, we first used 9-4c, a monoclonal antibody that visualizes the pallio-subpallial (PSP) boundary via recognition of reticulon 1-A and -B (Hirata et al., 2002). We found that the pallium (which is dorsal to the 9-4c^+^ band and consists of the dorsal, lateral, and ventral pallium [Puellus et al., 2000]) progressively expands ventrally during the embryonic period (Figure 1C), externally covering (“riding on”) the striatum by E13. Further analysis from E10 to E13, based on anti-Pax6 and anti-Tbr1 immunohistochemistry (Figure 1D), revealed that the “riding” of the striatum by the pallium (below the level of the PS angle) was accompanied by a disproportional (dorsal < ventral) thickening of an outer superficial zone containing Tbr1^+^ neurons (i.e., PP) in the pallial wall (portion dorsal to the PS angle).

During this early (∼E13) period, Nestin^+^ RFs in the pallial wall exhibited an oblique alignment pattern at the distal part (clearly seen at E12 and more prominent at E13) (Figure 1E). Careful inspection of the PSP boundary and the dorsally neighboring region (using the PS angle as a morphological reference) covered with Laminin^+^ basal lamina at E11 showed that the distal-most part of RC2^+^ RFs was ventrally deviated in a thin (one- or two-cell-thick) superficial zone (Figures 1F and S1A) that contained tangentially oriented PP neurons (βIII-tubulin^+^ and Tbr1^+^) (Figure S1B). This observation was supported by scanning electron microscopy (Figure S1C). These results led us to hypothesize that the neurons born in the dorsal pallium by E11, which constitute PP, might ventrally migrate to increase the horizontal size or area of a superficial tissue filled with pallial-derived cells—analogous to the manner in which the Nile Delta pushes out into the Mediterranean sea—thereby mechanically bending or deviating the distal part of RFs like wind bends trees (Figure 1G).

### Neurons born in the dorsal pallium at E10, including presumptive SP cells, migrate ventrally to stream PP

To determine whether E10 pallial-born neurons migrate ventrally, we used *in utero* electroporation (IUE), which is useful for labeling neurons that contribute to SP (Shimogori and Ogawa, 2008; Ohtaka-Maruyama et al., 2018). Live monitoring (at E11 or E12) of flat-mount cerebral walls (Figures 2A and 2B, Video S1A) and coronal slices (Figures 2C and 2D, Videos S1B and S1C) prepared following IUE at E10 revealed that heavily IUE-labeled cells constituted PP and underwent a massive ventral migration. This PP flow occurred beyond the level of the ventricular PS angle, expanding the pallial territory in the basal/superficial (neuronal) zone and curving the PSP boundary, which was confirmed *in vivo* (Figures S2A, S2B, and S2C). They exhibited locomotion-like salutatory behaviors, with average velocities of 32.2 μm/hr at E11 (n=20) and 21.1 μm/hr at E12 (n=18), sometimes changing direction via formation of a new leading process (Figure 2D, Video S1C). Ventral nucleosomal translocation of these neurons in the lateral region continued until E13 (Figure 1E, Video S2A) with a much smaller velocity (4.2 μm/hr, n=25), but was no longer observed at E14 (Figure 1F, Video S2B). Although a subset of the E10 pallial-born neurons (ventral-most fraction) seemed to contribute to the striatum and olfactory cortex, as previously suggested (Hamasaki et al., 2001; García-Moreno et al., 2008; Cocas et al., 2009) (Figures S2A and S2B), the majority of these cells born in the dorsal and dorsolateral pallium (judged so based on diluted but recognizable VZ labeling) were found in SP at E14 and later (Figures 2F, S2D-S2H), as determined by immunodetection of MAP2 (Crandal et al., 1986; Chun and Shatz, 1989) (Figures S2E and S2F) and Nurr1 (Hoerder-Suabedissen and Molnár, 2013) (Figure S2G).

**Figure 2.**
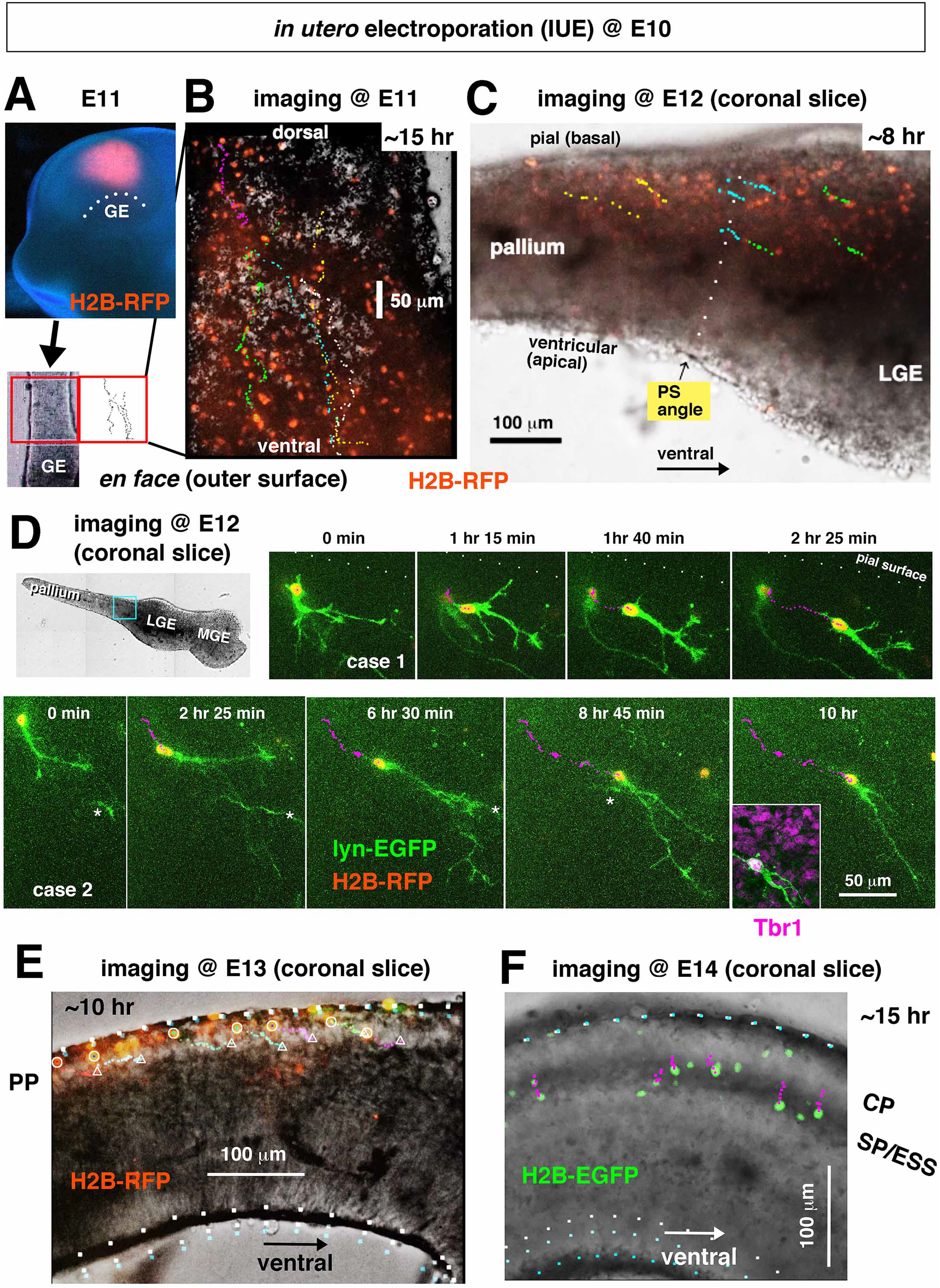
Ventral migration of E10 pallial-born neurons live-monitored until E13. (A) Low-power photomicrograph of an E11 hemisphere that received IUE at E10 in the pallium. GE, ganglionic eminence. (B) A cerebral wall flat-mount prepared from the hemisphere shown in (A), observed basal *en face* from E11 for 15 hr (Video S1A). Trajectories of neurons that migrated ventrally are superimposed. (C) A coronal slice monitored at E12 for 8 hr, following IUE at E10 (Video S1B). (D) High-resolution time-lapse monitoring of E10-born PP neurons in an E12 coronal slice, following IUE at E10 (Video S1C). (E) A coronal slice monitored at E13 for 10 hr (Video S2A), showing slower ventral displacements of E10-born neurons. (F) A coronal slice monitored from E14 for 15 hr (Video S2B), showing E10-born neurons contributing to SP/ESS (dark band) without ventral displacements. Note that while GFP^+^ nuclei of SP neurons translocate inward (towards the ventricular side), the apical/ventricular surface of the pallial wall (outlined) seems to be pushing outward.

Introduction of dominant-negative forms demonstrated the dependence of this ventral PP stream on Rac1 and N-Cadherin (Figure S2I), similar to the radial migration of glutamatergic neurons and ventral-to-dorsal migration of GABAergic interneurons (Kawauchi et al., 2003; 2010; Luccardini et al., 2013). Our immunohistochemical analysis did not reveal any abnormalities in the ventral extension of PP at E12 in *reeler* mice (data not shown) or in mice lacking CXCR4 (the receptor of CXCL12, which supports tangential migration of CR cells [Borell and Marin, 2006]) (data not shown).

### Ventrally locomoting E10-born neurons exert pulling and pushing forces and spatially interact with distal RFs

We next sought to assess the possible pushing effect of the PP stream to RFs, modeled by analogy to the pushing of trees by wind (Figure 1G). Because actomyosin is a major force generator in migrating neurons (Bellion et al., 2005; He et al., 2010; Trivedi et al., 2014) (Figure 3A), we performed immunohistochemistry with antibody against phosphorylated myosin light chain (p-MLC); the results revealed that PP neurons were positive for p-MLC (Figure 3B). To determine whether PP neurons pull at the tip of leading process, which should form the mechanical basis of pushing at the soma (Figure 3A), we took four different approaches. First, slice cultures showed that when a growth cone was extended by a PP neuron, a fluorescent spot in the surrounding space moved towards the growth cone (Figure 3C). Second, when explants of E11 or E12 pallial walls were cultured in soft collagen gel containing microbeads (Minegishi et al., 2018; Umeshima et al., 2018), the microbeads moved towards growth cones extended from the explants (Figure 3D, Video S3A). Third, when explants were cultured on plastic dishes, they crawled, and the front/leading portion was occupied by PP neurons (Figure S3A, Video S3B). Fourth, when coronal slices (freed from meninges) were cultured in soft collagen gel containing fluorescent microbeads, ventral-to-dorsal and slice-directed displacement of the beads was observed along the basal, but not apical, surface (Figure 3E, Video S3C). These results clearly show that PP neurons ventrally migrating along the basal/outer surface of the pallial wall exert pulling from their front. Initially (E11), the basal lamina is the major target for pulling by superficially migrating PP neurons. A little later (E12), pulling within PP may occur more deeply and intercellularly between migrating neurons in a manner that is shared by several types of collective cell migration (i.e., preceding cells are pulled by follower cells) (Bazellières et al., 2015; Vassilev et al., 2017), possibly involving N-cadherin (Figure S2I).

**Figure 3.**
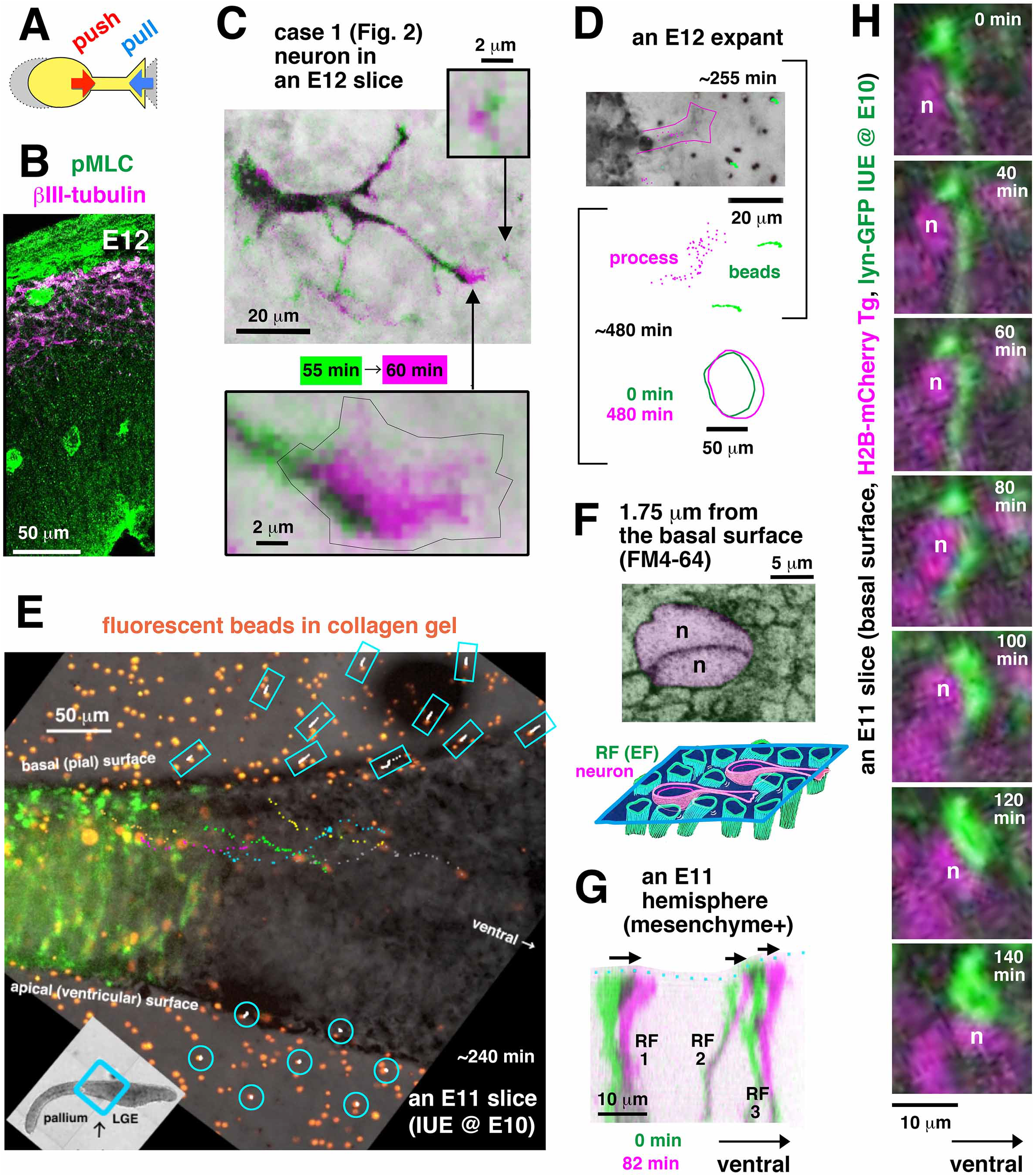
Pulling and pushing exerted by PP neurons until E12. (A) Schema of a locomotion-like migrating neuron, with expected pulling at the leading process and pushing at the soma. (B) E12 cerebral wall stained with anti–βIII-tubulin and anti–phospho-myosin light chain (pMLC), showing expression of pMLC in PP neurons. (C) Co-occurrence of the formation of a large growth cone at the tip of a leading process by a ventrally migrating neuron (case 1 in Figure 2) and displacement of a fluorescent spot in the surrounding space towards the growth cone. (D) Time-lapse monitoring of an E12 explant prepared from the PP region and embedded in soft collagen gel containing microbeads (Video S3A). Tracking of a single process and two beads at a high magnification (∼255 min in photomicrograph; ∼480 min summarized below) and that of the whole explant (circled at lower magnification, ∼480 min) show that advancement of the process and the explant coincided with pulling of the beads. (E) An E11 coronal slice cultured in soft collagen gel containing fluorescent microbeads. Monitoring for 240 min at the region near the PSP boundary showed that beads embedded in the outer (basal/pial) side of the slice moved towards the slice and dorsally, whereas those in the opposite (apical/ventricular) side did not move (Video S3C). Some E10-born neurons were tracked to show their ventral migration. (F) Confocal *en face* observation of an FM4-64–stained E12 pallial wall from the outer surface (Video S4A for 3D assessment). Mesh-like RF endfeet and neurons (n) coexist on the outer surface. (G) High-resolution live monitoring of three RFs in an E11 pallium covered with intact meninges (mesenchymal sheet) (Video S4B). (H) Simultaneous monitoring of a single RF (GFP-labeled via IUE) and nuclei of PP neurons in a pallial wall prepared from an E11 H2B-mCherry transgenic mouse (Video S4C).

We next asked whether, and if so how, such pulling is associated with pushing (of neighboring cellular structures, including RFs) by the soma. Confocal microscopy of FM4.64-stained pallial walls revealed that the basal surface of the wall consisted of dense RF endfeet and neurons (Figure 3F, Video S4A). This observation, together with a similar electron microscopy finding (Shoukimas and Hinds, 1978), is consistent with the idea that moving PP neurons geometrically compete with RF endfeet for the basal lamina. We then time-lapse monitored RFs in an E11 pallial wall covered closely with meninges, and found that they gradually deviated ventrally, with the endfeet translocating ventrally (Figure 3G, Video S4B). Dual-color monitoring at or near the basal surface revealed that when the nuclei and somata of neurons moved ventrally, RF endfeet also translocated ventrally (Figures 3H and S3B, Videos S4C and S4D). Thus, the multicellular dynamics in PP, which orthogonally contains RFs, is consistent with a model in which the PP stream participates in realignment of RFs on and near the basal surface (Figure S3C).

Culturing slices in bead-containing soft gel, which can be easily deformed, is most suitable for detecting pulling that occurs at the outer surface of slices (Figure 3E), but is not ideal for observing pushing in that region. To determine whether the meningeal mesenchyme, which is enriched with p-MLC (Figure 3B) and in tight contact with PP neurons (Figure 1F) *in vivo*, can resist (like the dish surface) pulling by PP neurons (at the leading process) and thus counteract it, thereby allowing the PP neurons to elicit a pushing force at the soma (Figure S3C), we assessed its mechanical properties. When embryos at E10–12 were treated under hypoxic conditions for 1–2 hr (by placing excised uteri into culture medium and incubating them in a CO_2_ incubator), the cranial soft tissue composed of the meningeal mesenchyme and the overlying epidermis shrank, making the inner cerebral wall buckle (Figures S3D and S3E). Because the meningeal mesenchyme contains abundant vascular plexuses, this can be explained in part by hypoxia-induced contraction of peripheral vessels (Brinks et al., 2016). This experimentally revealed (over-induced) tangential contractility of the brain-encapsulating meningeal mesenchyme was further confirmed as follows. E11 or E12 cranial tissues were isolated from the cerebral wall based on differential (shrinking vs. buckling) responses to hypoxia (Figures S3D and S3E); otherwise, the meningeal/mesenchymal tissue and the cerebral wall stick to each other. When these tissues were directly pulled and released, they recoiled to shrink (Figure S3F). These results suggest that the meningeal mesenchyme may normally be under tangential tension, mechanically providing anchoring points to the pulling PP neurons, thereby aiding the soma-mediated pushing by PP neurons that occurs superficially in the pallial wall at E11.

At E12, pushing by the soma can clearly be inferred in a deeper zone where some PP neurons are migrating in the lateral pallium; RFs were displaced ventrally where PP neurons passed through (Figure S4A). In such lateral regions, the dorsal-to-ventral stream of PP was sandwiched by two zones where ventral-to-dorsal streams of interneurons (De Carlos et al., 1996; Anderson et al., 1997; Tamamaki et al., 1997) and glutamatergic neurons (Teissier et al., 2010) (Figure S4B) use, not colliding with them.

### PP neurons during the transition into the CP stage are morphologically and behaviorally heterogeneous

During the transition from the PP stage (with maximal ventral streaming, ∼E12) to the CP-forming stage (lacking ventral streaming, ∼E14), the site of obliqueness that occurred in RFs became farther from the basal surface (Figure 4A). Just before and during this transitional step (mainly at E13), corticofugal axons start to grow (Kim et al, 1991; De Carlos and O’Leary, 1992; Erzurumlu and Jhaveri, 1992; Molnar et al., 1998; Auladell et al., 2000; Espinosa et al., 2009). Hence, we asked whether, and if so how, pioneering axons participate in this transitional PP stream (which is slower than the earlier stream, Figure 2). Live inspection at E13 (and E14, if dorsal), following IUE-based labeling at E10 reveled that 61% of the total E10-born neurons observed (n=44) did not have axon-like long processes, and that their nucleosomal movements were preceded by not only ventral but also inward extension of short (20–30 μm) processes (Figure 4B, Video S5A). The remaining population (39% in the same E10 IUE-based counting) had long axons running beyond the PSP boundary (Figure S4C). Such axon-bearing PP neurons were also visualized by retrograde labeling with DiI from the internal capsule (Figures 4C and 4D). The nucleus of each axon-bearing PP neuron moved into the axon running ventrally and inward (Figures 4C and 4D).

**Figure 4.**
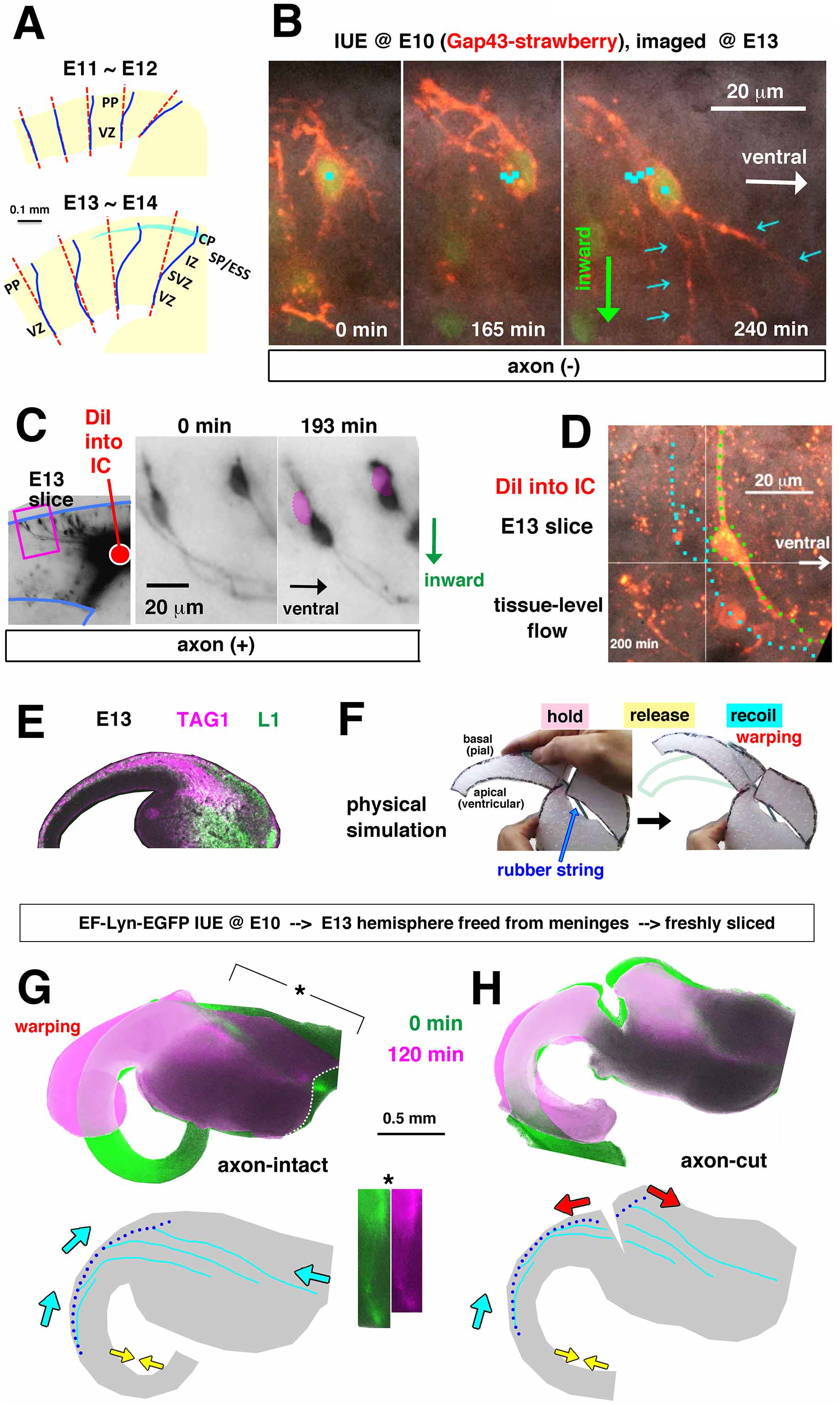
Behaviors of PP neurons, including axon-bearing neurons, in E13 pallial walls. (A) Schematic illustration summarizing deflection of RFs, which is deeper at E13 than at earlier stages. (B) Monitoring of a single E10-born neuron in an E13 coronal slice, with tracking of non-axon processes (small arrow) and the nucleus (Movie S5A). (C, D) Live observation of PP neurons retrogradely labeled with DiI inserted into the internal capsule (IC) at E13 (Movie S5B for D). (E) Anti-TAG1 (magenta) and anti-L1 (green) immunostaining at E13. (F) Physical model simulating a possible pulling force by axons extended corticofugally to the subpallium. The elastic property given to the axon by a rubber string causes a warping response of the pallial wall (Movie S5C). (G, H) Photomicrographs of coronal cerebral slices (meninges-free) prepared at E13. Merged bright-field and GFP images of a control slice (G) and an axon-cut slice (H) taken at 0 min (green) and 12 min (magenta) are shown. The intact slice (G) exhibited an outward warping of the pallial wall (cyan rightward arrows) (see Movie S5D for another example) and shrinkage of the ventral edge of the subpallial part (cyan leftward arrow), with a shortening of the EGFP^+^ corticofugal axons (asterisk, light blue illustrated below) from the E10-born PP/SP neurons (blue dots). The axon-cut slice (H) almost entirely lost the extensive warping response, and instead exhibited bidirectional shrinkage (red arrows) of the cut edges. Narrowing of the apical/ventricular surface (yellow centripetal arrows) occurred in both slices.

These axon^+^ neurons enabled us to detect collective displacement of PP neurons (Figure 4D, Video S5B), in which all neuronal elements (axon, soma, and dendrite-like outward-extended process) together moved exclusively ventrally, independently of the cell-intrinsic, intra-axonal nucleokinesis. RFs live-monitored in the lateral region of E13 slices exhibited an abrupt bending at the level of presumptive SP (interface between PP and the intermediate zone), where axons from more dorsal regions grow ventrally to form the external sagittal stratum (ESS) (Misson et al., 1988) (Figure S4D, Video S6A). Because axons exhibit traction forces (Bray, 1984; Lamoureux et al., 1989) and corticofugal pioneering axons have larger growth cones than follower axons (Kim et al., 1991), it is possible that axons ventrally growing (Figure 4E) participate in the streaming of PP via traction (Figure S4E). Our physical model (Figure 4F, Video S5C) suggested that if pallial walls containing an elastic rubber string (representing axons in tension) are released from a holding/constriction by the surrounding meningeal and epidermal tissues (Figures S3D, S3E and S4F), it would recoil outward, exhibiting a “warping” response, like the opening of the pine cone scales (Dawson et al., 1997).

Indeed, when E13 cerebral walls were isolated from the skull and freed from the meninges (and thus released from intracranial mechanical restriction *in vivo*) and quickly sliced coronally, keeping the pallium and subpallium fully connected, the slices exhibited outward warping of the pallial wall (Figure 4G, Video S5D) (n=5/5). This warping was quantitatively characterized by a reduction in the distance from the intra-pallial PP to the ventral edge of the subpallium (to 91±3%), between which the corticofugal axons from PP neurons run (Figure 4G), reproducing the results of the physical simulation. The warping occurred a little later (60–120 min) than the inward curling of the pallial wall (∼40 min), which is dependent on actomyosin along the apical/ventricular surface (Okamoto et al, 2013; Shinoda et al., 2018). When we made surgical incisions crossing the corticofugal axons (Figure 4H), the slices exhibited much weaker warping (n=5/5), with shrinkage of the cut edges in the dorsal and ventral directions, as well as significantly milder shortening (to 96±2%, p=0.024 [Exact Wilcoxon rank–sum test]) of the distance from the intra-pallial PP to the ventral edge of the subpallium. Together, these physical examinations support the *in vivo* occurrence of traction by corticofugal axons derived from PP neurons. Thus, morphologically and behaviorally heterogeneous PP neurons may collectively stream ventrally from E11 to E13 and interact mechanically with RFs in a stage-dependent manner (Figure S4G).

### Experimental weakening of PP stream decreases RF deviation by E13

To determine whether the PP stream is necessary for the ventral curving/deflection of RFs, we decreased the number of E10 pallial-born neurons via IUE-mediated expression of the diphtheria toxin A chain (DTA) under the control of the *NeuroD* promoter (NeuroD-DTA), which killed differentiating cells as early as 12 hr after IUE without harming undifferentiated RF-forming progenitors (radial glial cells, RGCs) (Watanabe et al., 2018) (Figure S5A). Neuro-DTA IUE at E10 did not affect the spatial expression patterns of FGF8, Wnt3a, Emx1, or Pax6 at E11 (Figure S5B), and we observed no excessive cell death in VZ (Figure S5C). Neurons born later than E10, e.g., Ctip2^+^ neurons (Figure S5D) or those labeled with bromodeoxyuridine (BrdU) administered at E12 (Figure S5E), did not die excessively. We quantitatively assessed the morphology of RGCs labeled from the outer pallial surface with DiI (Miyata et al., 2001) (Figures 5A, 5B and 5C). We compared the PP-ablated hemispheres (n=26 RGCs) and normal (non-IUE, contralateral) hemispheres (n=48 RGCs) at E13 using two different approaches (length ratio and angle; see Methods). We found that the ventral deflection of RGCs from the shortest ventricle-to-pia line was seen much more clearly in the lateral part (closest to the PS angle) making a crease/angle at the level of presumptive SP (or ESS) than in the dorsal region (far from the PS angle) (R^2^ = 0.44 and 0.31 for the length ratio and angle, respectively). We then found that RGC deflection was much less evident when the PP stream was weakened (R^2^ = 0.05 and 0.02) (p=0.02 and 0.04, ANCOVA) (Figures 5D, 5E, and S5F). These results suggest that the PP stream contributes to the RF deviation/deflection that occurs by E13.

**Figure 5.**
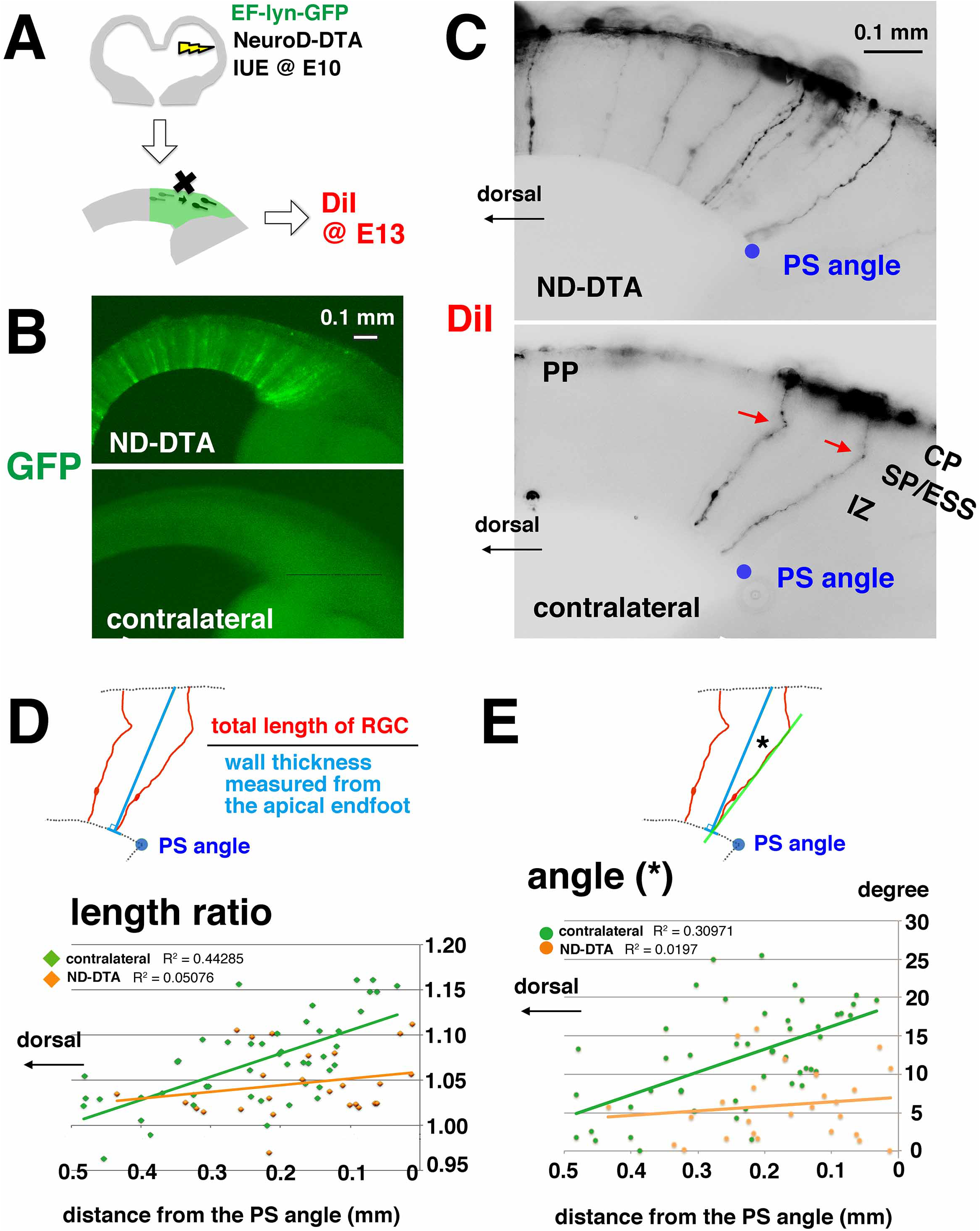
Weakening of PP stream results in smaller deflection of RFs by E13. (A, B) Experimental design. IUE with EF-lyn-GFP with NeuroD-DTA at E10 was followed by DiI labeling of the outer surface of pallial walls at E13. (C) Representative DiI-labeled RGCs in a NeuroD-DTA–electroporated (also GFP-labeled) hemisphere and those in the contralateral hemisphere. (D) Comparison of the length ratio (total length of each RGC divided by the thickness of the wall measured from the apical endfoot). Wall thickness was measured by drawing a line (blue) perpendicular to the surface. R^2^=0.44 for contralateral or non-IUE (n=48 RGCs from eight embryos), and R^2^=0.05 for NeuroD-DTA (n=26 RGCs from four embryos) (p=0.02, ANCOVA). (E) Comparison of the angle of RF deflection (designated as an angle formed by the inner portion of each RGC and the shortest line from the apical endfoot of that RGC). R^2^=0.31 for contralateral or non-IUE, and R^2^=0.02 for NeuroD-DTA (p=0.04, ANCOVA).

### Deep-layer neurons obliquely enter PP to initiate CP

Inspection of GFP-labeled neurons at E15, following IUE to the dorsal region at E10 (Figure S6A), showed that weakly labeled cells corresponding to neurons destined for layer 6 (in addition to the most heavily labeled SP-like cells) spread laterally, consistent with a previous clonal analysis (Gao et al., 2014). Their ventral spreading was less extensive than that of heavily labeled E10-born neurons, but was beyond the extent of the GFP^+^ cells in VZ (Figures S6A, S6B, and S6C). To determine how CP is initiated by these laterally spreading deep-layer neurons in the presence of the PP stream, we live-monitored pallial walls in which the nuclei of all cells, including CP-initiating cells, could be visualized using the H2B-mCherry transgene (Abe et al., 2011). We found that the entrance of neurons into a newly emerging CP at E14, as well as their intra-CP migration, were frequently oblique (ventrally deviated) (Figure 6A, Movie S6B). At E15, however, new neurons entered and migrated in CP almost perpendicularly towards the pial surface (Figure 6B). The oblique neuronal entrance observed at E14 could be explained by the obliqueness of RFs relative to the guide neurons. Additionally, the PP stream might drag the PP-entering (CP-initiating) neurons ventrally (Figure S6D). Both models are consistent with the possibility that the initiation and the subsequent growth of CP are assisted by the preexisting PP stream. We also found that CP initiation by the entering neurons was accompanied by inward somal translocation, as well as reorientation (radial _➔_ horizontal) of SP-like neurons (Figure S6E).

**Figure 6.**
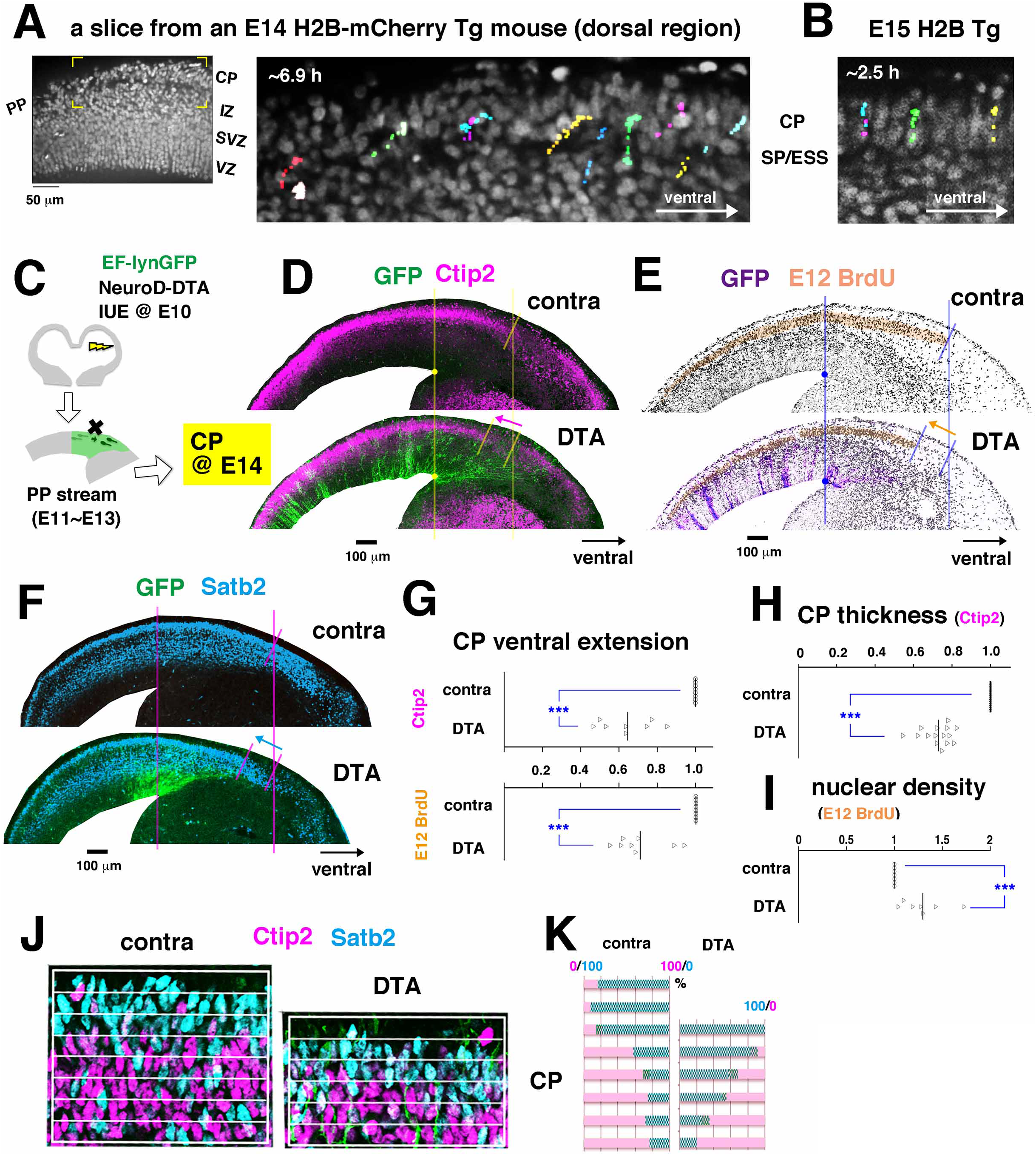
CP initiation by the entrance of deep-layer neurons under the influence of ventrally streaming PP. (A) A coronal slice prepared from an E14 H2B-mCherry mouse, monitored for 6.9 hr (Video S6B). (B) A coronal slice prepared from an E15 H2B-mCherry mouse, monitored for 2.5 hr. (C) Experimental design. IUE with EF-lyn-GFP with NeuroD-DTA at E10 was followed by BrdU pulse-chase at E12 and analysis at E14. (D) Anti-Ctip2 (magenta) and anti-GFP (green) immunostaining. (E) Anti-BrdU (blue) and anti-GFP (purple) immunostaining. CP is orange highlighted. (F) Anti-Satb2 (blue) and anti-GFP (green) immunostaining. (G) Comparison of the length of CP extended ventrally beyond the line drawn from the PS angle (as in D and E) between the DTA-IUE and the control (contralateral) groups. Each graph depicts the ratio, with the value for the control regarded as 1.0 (p=0.0002 for anti-Ctip2 analysis [n=8 sections from three embryos]; p=0.0002 for anti-BrdU analysis [n=8 sections from three embryos]). (H) Comparison of the thickness of CP extended ventrally beyond the line drawn from the PS angle (as in D) between the DTA-IUE and the control (contralateral) groups. Graph depicts the ratio, with the value for the control defined as 1.0 (p=1.3×10^−8^, n=15 regions in six sections from three embryos). (I) Comparison of the density of BrdU^+^ nuclei in CP extended ventrally beyond the line drawn from the PS angle (as in E) between the DTA-IUE and the control (contralateral) groups. Graph depicts the ratio, with the value for the control defined as 1.0 (p=0.0006, n=7 sections from three embryos). (J) Anti-Ctip2 (magenta) and anti-Satb2 (cyan) immunostaining. (K) Graph depicting the proportions of Ctip2^+^ or Satb2^+^ nuclei in each bin (10 μm) of the ventralmost CP in a representative section.

### Experimental weakening of the PP stream decreases ventral extension of CP

To investigate the aforementioned functional contribution of the PP stream to CP formation/growth, we electroporated embryos with NeuroD-DTA at E10, administered BrdU at E12, and then performed immunohistochemistry for BrdU, Ctip2, and Satb2, at E14 (Figure 6C). We focused on the lateral (ventralmost) CP region into which neurons originating from the most ventral pallial VZ (closest to the PS angle) could migrate (Figures S6B and S6C). The lateral CP was less ventrally extended in the DTA-electroporated walls than in the contralateral walls (Figures 6D–G); note that CP extension was indistinguishable between the GFP-electroporated wall (i.e., the true control) and the contralateral un-electroporated wall (Figure S6C). The less ventrally extended (p<0.001) CP in the PP-reduced (NeuroD-DTA-E10–electroporated) hemispheres was thinner (Figure 6H) (p<0.001) and contained a higher density of BrdU^+^ nuclei (Figure 6I) (p<0.001) than normal CP (exact Wilcoxon rank–sum test). Although the inside-out segregation pattern of early-born (Ctip2^+^, deep) neurons and later-born (Satb2^+^, relatively superficial) neurons was conserved in these abnormal CPs formed under compromised PP streaming, it was vertically compressed (Figures 6J and 6K). These data strongly suggest that PP streaming until E13 assists in the ventral-ward formation of CP.

### Experimental weakening of the PP stream dorsally shifts the postnatal neocortical map including somatosensory barrels

To determine whether the early embryonic PP stream is necessary for the normal and ordered establishment of functional mapping in the postnatal neocortex, we subjected E10 Ai14 mice to *in utero* electroporation with NeuroD-DTA or control CAG-GFP vectors together with EF-cre, which allowed us to retrospectively identify the occurrence and position of IUE at E10 through postnatal detection of TdTomato fluorescence (Figures 7A and S7A). TdTomato^+^ hemispheres at postnatal day 8 (P8) were subjected to *in situ* hybridization (ISH) for Rorβ (in coronal sections) and cytochrome oxidase (CO) in flat-mount sections (Figure 7A). Comparison between the electroporated side and the contralateral side in CAG-GFP–electroporated hemispheres showed that the arealization pattern visualized with Rorβ ISH was completely symmetrical, irrespective of the site of IUE (TdTomato^+^ region) (n=3/3) (Figure 7B). By contrast, the same right–left comparison for NeuroD-DTA–electroporated hemispheres revealed dorsal shifts of Rorβ^+^ domains on the IUE side (Figure 7C). In hemispheres that were electroporated in the lateral region, lateral Rorβ^+^ domains such as Au1 or S2 shifted dorsally (Figure 7C, 1 and 3). When IUE was limited to the lateral region, domains located much more dorsally (such as S1) were not affected (Figure 7C, 1). In hemispheres that were only dorsomedially electroporated, S1 labeling with Rorβ probe (Figure 7C, 4) or anti-Cux1 (Figure S7B) shifted dorsally. In addition, wider IUE involving both the dorsal and lateral regions, including the intermediate dorsolateral region, resulted in dorsal shift of multiple Rorβ^+^ regions (Figure 7C, 1, 2, and 3). These results can be summarized as a rule (n=6/6): regions that should have been formed downstream of the IUE position shifted dorsally in the NeuroD-DTA group.

**Figure 7.**
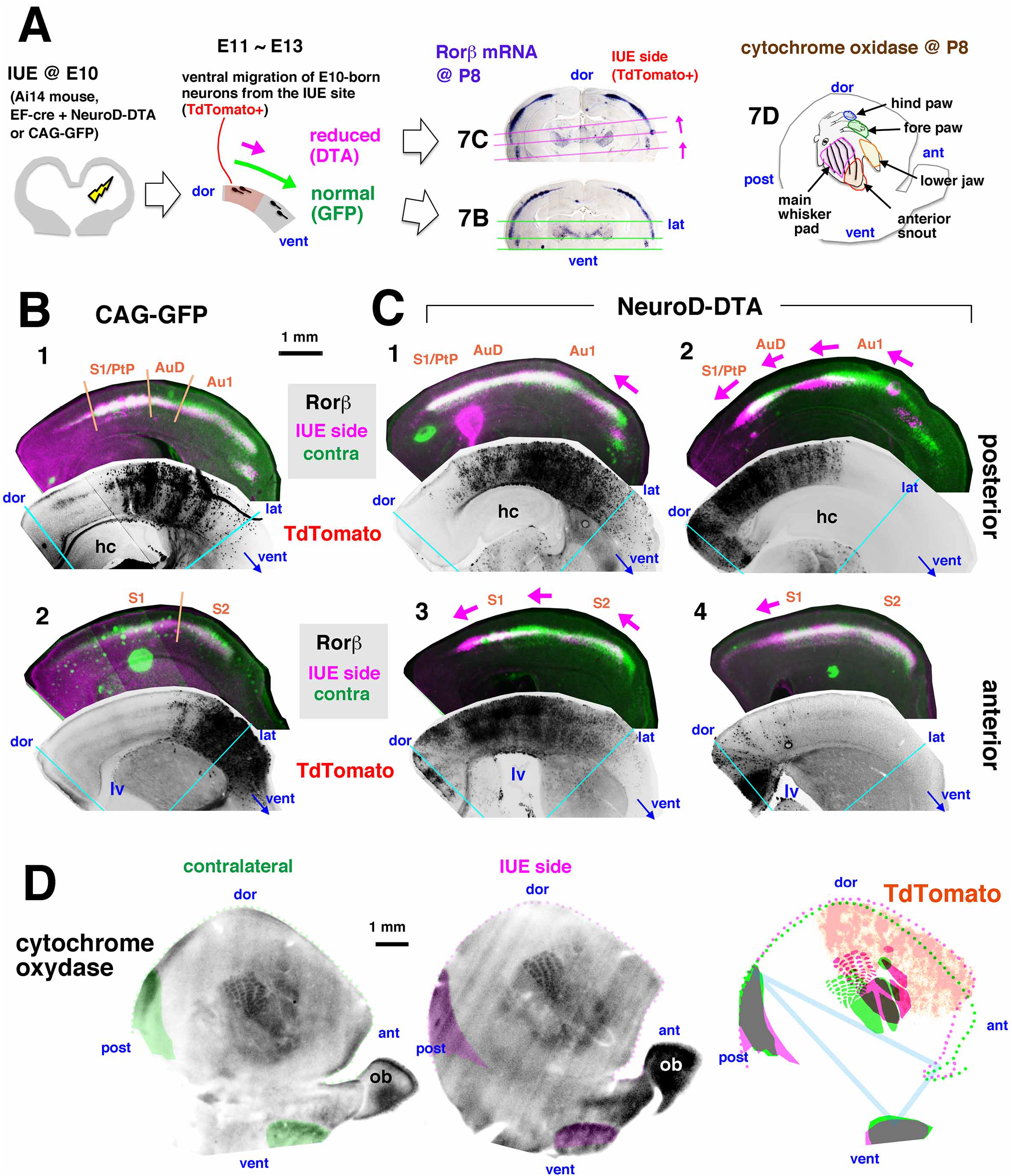
Early embryonic depletion of PP streams resulted in dorsal shifts of the postnatal neocortical maps. (A) Experimental design. Ai14 mice were electroporated at E10 with EF-cre and CAG-GFP or NeuroD-DTA, followed by *in situ* hybridization for Rorβ and cytochrome oxidase stain at P8. (B) Left–right comparison of Rorβ expression between the CAG-GFP-IUE side (magenta) and the contralateral/non-IUE side (green), showing almost identical/symmetrical patterns, regardless of the position of IUE visualized with TdTomato. hc, hippocampus. lv, lateral ventricle. According to Barber et al. (2015): S1, primary somatosensory cortex; S2, secondary somatosensory cortex; S1/PtP, primary sensory/parietal posterior association cortex posterior rostral and dorsal; AuD, auditory secondary cortex dorsal; Au1, primary auditory cortex. (C) Left–right comparison of Rorβ expression between the NeuroD-DTA-IUE side (magenta) and the contralateral/non-IUE side (green), showing dorsal shifts (arrow) of Rorβ^+^ domains that were to be formed downstream of the origin of the ablated PP neurons (i.e., ventral flow), faithfully reflecting the position of IUE visualized with TdTomato. (D) Comparison of cytochrome oxidase staining between the NeuroD-DTA and contralateral sides.

This apparent rule also applied to CO-stained hemispheres (n=3/3): compared to CO^+^ barrels in the contralateral (non-electroporated) hemispheres, those in NeuroD-DTA–introduced hemispheres shifted dorsally (i.e., towards the TdTomato^+^ area) (Figures 7D, S7C, and S7D). The dorsal shift of somatosensory areas that we obtained by decreasing the number of E10 pallial-born PP neurons without killing CR neurons was similar, in part, to the results obtained in Barber et al. (2015), who disturbed the normal embryonic tri-territorial distribution of CR cells.

Some barrels, as well as the total posteromedial barrel subfield (PMBSF) (Moreno-Juan et al., 2017), were a little larger on the contralateral side (Figures S7C and S7D), consistent with the involvement of SP neurons in barrel formation (Piñon et al., 2009; Hoerder-Suabedissen and Molnár, 2015). In summary, neurons born in the pallium at E10 that contain presumptive SP neurons and comprise the PP ventrally stream until E13 to mechanically support the initiation and growth of CP, thereby contributing to the normal formation of postnatal somatotopic fields in the neocortex.

## Discussion

### PP neurons play frontier-advancing and pre-patterning roles in the early embryonic pallium

Rearrangement or repositioning of cells is a fundamental mechanism by which a developing organ acquires a functional structure. For example, tissue-level assembly of cellular movements in which two cells that were originally aligned along the left–right axis exchange their neighboring cells (i.e., intercalate) to become aligned along the anterior–posterior (AP) axis, enables a growing neural plate to elongate along the AP axis (to eventually form the brain and spinal cord). Likewise, the neocortex and the striatum, which were in a dorsal–ventral relationship during the early embryonic primordial period, acquired a superficial–deep relationship in the postnatal/adult mammalian cranium. This rearrangement, which couples an inward growth of the striatum to narrow the lateral ventricle and a ventral spreading of the neocortex to externally or superficially cover the striatum (to be worn as a “pallium” [= vestment] by the striatum), promotes space efficiency in the fetal cranium, whose growth is ultimately constrained by the size of the birth canal. Recent studies revealed that embryonic brains that develop under space limitations are sensitive to the physical conditions faced by their cells during development (Okamoto et al., 2013), and that developing brains utilize physical or mechanical factors that emerge as a function of the morphology, 3D assembly pattern, and behavior of brain-forming cells (Koser et al., 2016; Karzbrun et al., 2018; Long et al., 2018; Shinoda et al., 2018; Watanabe et al., 2018; reviewed in Franze, 2013; Llinares-Benadero and Borrell, 2019). In this study, we identified another way in which cells use an externally provided force to fulfill their developmental roles. The asymmetric (ventralward) deflection of RFs, which forms the basis for the protomap mechanism (Rakic, 1988) underlying the establishment of functional neocortical maps, depends on the ventral migration of PP neurons (including future SP neurons) born early in the pallium.

In addition to their previously known roles in layer formation (Ogawa et al., 1995; Ohtaka-Maruyama et al., 2018) and the connection between the cortex and thalamus (McConnell et al., 1989; Ghosh et al., 1990; reviewed in Allendoerfer and Shatz, 1994; Hoerder-Suabedissen and Molnár, 2015), PP neurons promote, through their ventral streaming via tangential migration (E11–12) and axon-based traction (E13), the ventral extension of CP (E14), thereby playing a priming physical role in postnatal neocortical arealization. A recent study by Barber et al. (2015) showed that appropriate and balanced tangential migration of CR cells from three different origins (i.e., the hem, septum, and PS angle) to cooperatively cover the entire pallial surface by E12 underlies the normal regionalization of postnatal neocortex. It remained unclear, however, how CR cells play this early embryonic “pre-patterning” role for postnatal arealization. Our discovery of ventral PP streaming and its contribution to CP formation and postnatal arealization supports the idea that CR cells are the first flow generators in the developing pallium, and that their movements influence physical behaviors of the non-CR PP neurons (as we observed in this study), thereby affecting the initiation and formation of CP. It is also possible that neurons generated around E11 and E12 that initiate CP (i.e., those comprising future layers 6 or 5) collaborate with the pre-existing PP neurons to pull at their corticofugally growing axons, thereby increasing the total (PP plus early CP) ventral flow of somata.

### Possible molecular mechanism for the ventral PP streaming

Rac1 and N-Cad seem to be important for the motility of PP neurons (Figure S2I). One candidate that could explain the dorsal-to-ventral directionality of the PP stream is Netrin 1. We detected a dorsal-to-ventral gradient of anti–Netrin-1 immunoreactivity in the E11 cerebral wall that increases from the pallium to the striatum (K. Saito, unpublished observation). Netrin-1 is implicated in the ventral migration of lot (lateral olfactory tract) cells from the dorsal pallium (Kawasaki et al., 2006), and it is also involved in the formation of corticofugal axons (Métin et al., 1997). PP neurons, including nascent SP neurons, express DCC (Kawasaki et al., 2006). Another candidate is semaphorin system (Bagnard et al., 1998), which controls corticofugal axonogenesis. Whether these molecular mechanisms underlie ventral PP streaming should be determined in future studies.

### Give-and-take of mechanical stresses in multicellular brain development

From E11 to E12, the basal/outer endfeet of RFs are ventrally displaced along the basal lamina (meninges), deflecting the most distal portion of RFs (≤ 30 μm from the surface) ventrally. This could simply occur passively, as in the endfeet of chicken retinal neuroepithelial cells co-cultured with tangentially migrating cells (Halfter et al., 1988), but may also involve RFs’ active crawling on the basal lamina, as suggested in the zebrafish retina (Sidhaye and Norden, 2017). In either case, pushing forces generated by the ventralward flow of the somata of PP neurons (based on traction at their leading processes) seem to be received by RGCs. At E13, similar soma-mediated pushing continues ventralward, but deeper, with a contribution of corticofugal axons to more long-range tractions. At E14, SP neurons, the major derivatives of PP neurons, no longer move ventrally, and instead gradually translocate inward (towards the ventricular side) (Figure 2F). Inward translocation of SP neurons is often led by short anchoring processes (Figure 4B) and coincides with another inward movement exhibited by the entire apical/ventricular surface consisting of RGC endfeet (Figure 2F). This apparent coupling raises the possibility that the settlement or anchoring of SP neurons that are to be passed through (rather than displaced outward) by newly coming layer 6 neurons might be aided by RGCs, possibly through anchoring of the short processes of SP neurons to RFs. This mechanism should be investigated in future studies.

Mutual ‘give and take’ of mechanical stresses (including elastic energy) between cells of different cell-cycle or differentiation status occurs in the VZ/SVZ; these processes streamline interkinetic nuclear migration (Shinoda et al., 2018; Watanabe et al., 2018), thereby assisting in cell production. This efficient tissue formation via a combination of active and passive movements is reminiscent of the minimization of total energetic expenditure in diverse biological events within ecosystems, e.g., birds in V formation, which fly alternatingly either as a leader to generate wingtip vortices or as followers to utilize a lifting force generated by the leader-driven vortices (Bajec and Heppner, 2009). The new, externally arising physical mechanism that we identified between PP neurons and RFs in the early corticogenic stage collaborates with previously elucidated cell-intrinsic (e.g., Emx2/Pax6 [Bishop et al., 2000]) and chemically mediated mechanisms (e.g., FGF8 [Fukuchi-Shimogori et al., 2001]) to fully explain how neocortical arealization is established.

## Legends for Supplemental Figures

**Figure S1.**
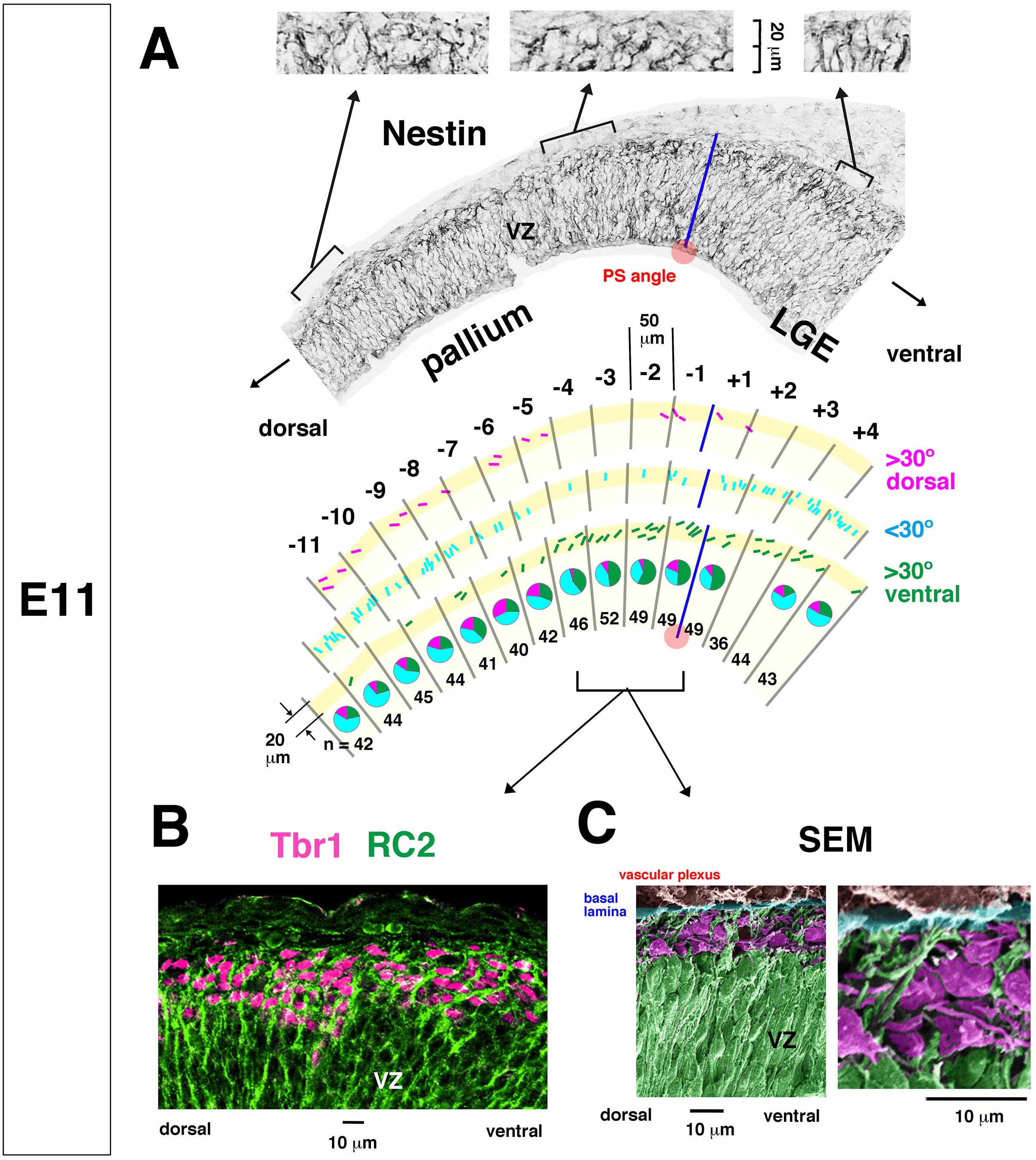
Ventral deflection of RFs detected superficially near the PSP boundary at E11, Related to Figure 1. (A) Anti-Nestin–immunostained sections were analyzed for the morphology of RFs in the superficial zone (within 20 μm from the pial surface). The traced distal RFs (exemplifying the results obtained from one section) were categorized into three groups (>30° deviated dorsally, >30° deviated ventrally, or intermediate [radially oriented]) in each 50 μm–wide column along the dorsal-to-ventral axis. Pie graphs summarize the results collected from three sections. Red spot, PS angle. (B) RC2 and anti-Tbr1 photomicrograph showing the coexistence of horizontally oriented Tbr1^+^ neuronal nuclei in the superficial zone containing ventrally deflected RFs in regions near the PSP boundary. (C) Scanning electron microscopy on an E11 pallial wall near the PSP boundary.

**Figure S2.**
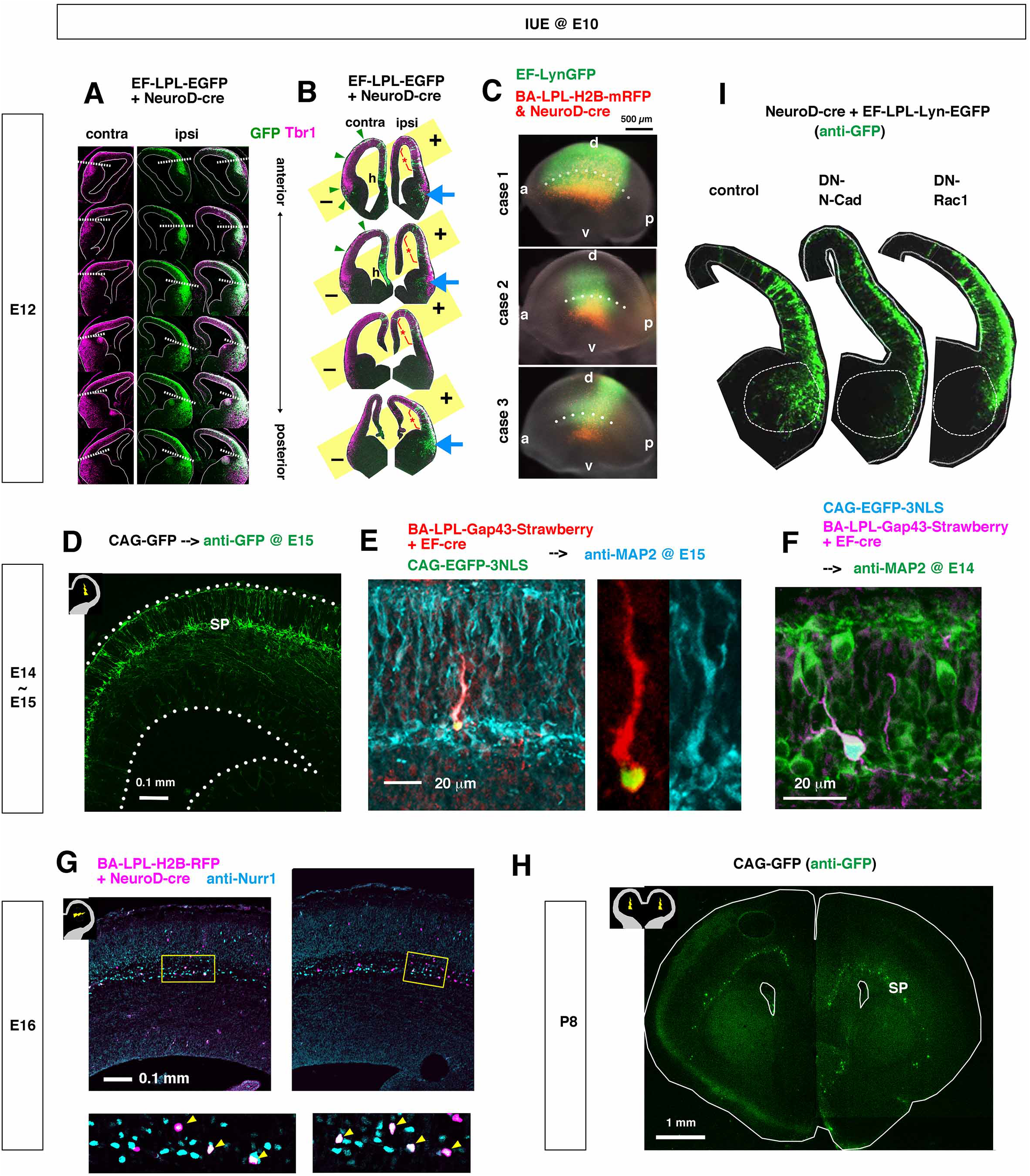
*In vivo* analysis of ventral migration of the E10 pallial-born neurons, Related to Figure 2. (A) Anti-GFP and anti-Tbr1 immunostained coronal sections of an E12 cerebrum that received unilateral intra-pallial IUE at E10. (B) Anti-GFP and anti-Tbr1 immunostained coronal sections of an E12 cerebrum that received IUE at E10. In the hemisphere whose dorsolateral pallium had been electroporated (asterisk, ipsi), neurons that migrated tangentially (thick arrow) were abundant in regions ventral to the PS angle. In the contralateral hemisphere that was electroporated in the dorsomedial region and the hem (h), a few PP neurons including Cajal–Retzius neurons also exhibited ventral dispersion (arrowhead). (C) Whole-mount fluorescent photomicrographs showing that neurons heavily (and almost specifically) labeled with RFP via IUE at E10 using NeuroD-cre were found more ventrally than the main IUE region, which was brightly GFP^+^ due to more continuous and progenitor-directed expression of EF-lyn-GFP. a, anterior; p, posterior; d, dorsal; v, ventral. (D) Anti-GFP photomicrograph showing that neurons labeled with IUE at E10 with CAG-EGFP constituted the subplate (SP) at E15, and had horizontally oriented morphologies. (E, F) Anti-MAP2 photomicrograph showing that a neuron labeled with IUE at E10 was found in the subplate (SP) at E14 (F) and E15 (E). (G) Anti-Nurr1 photomicrograph at E16 showing that neurons labeled with IUE at E10 were positive for Nurr1 (arrow). (H) Anti-GFP photomicrograph showing that neurons labeled with IUE at E10 (with CAG-EGFP) constituted the subplate (SP) at postnatal day (P) 8. (I) Comparison of the ventral spreading of E10 pallial-born neurons in E12 hemispheres. Note the lack of entrance of GFP^+^ neurons into the striatal region in hemispheres electroporated with dominant-negative forms of N-Cadherin or Rac1.

**Figure S3.**
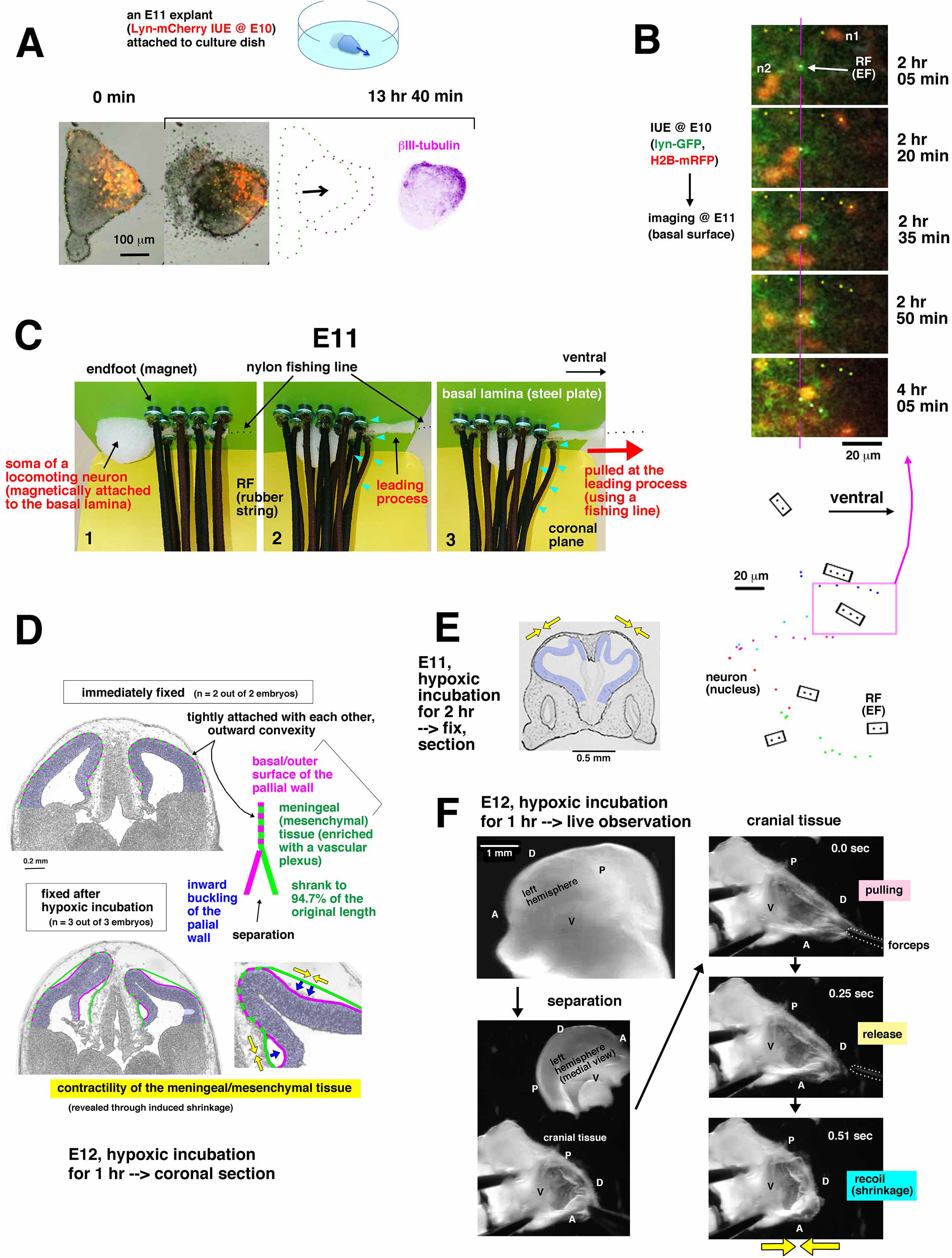
Mechanical assessment on PP neurons, RFs, and the surrounding connective tissue, Related to Figure 3. (A) An explant prepared from an E11 pallium that received IUE at E10 was cultured on a plastic dish coated with polyethylenimine (Video S3B), followed by anti–βΙΙΙ-tubulin immunostaining. (B) *En face* monitoring of the outer/basal surface of an E11 pallium that was electroporated at E10 (Video S4D). Endfeet (EF) of RFs (green) and neuronal nuclei (red) were tracked. (C) Physical model showing that a neuron migrating along the surrounding mesenchyme (based on its attachment to the basal lamina) would push the distal part of RFs via its soma. (D, E) Experimentally revealed (over-induced) tangential contractility of brain-encapsulating meningeal mesenchyme. When embryos at E10–12 were subjected to hypoxic condition for 1–2 hr (by placing excised uteri into culture medium and incubating them in a CO_2_ incubator), the cranial soft tissue consisting of the meningeal mesenchyme and the overlying epidermis shrank, making the inner cerebral wall buckle; this observation can be partly explained by hypoxia-induced contraction of peripheral vessels (Brinks et al., 2016), because the meningeal mesenchyme contains abundant vascular plexuses. (F) Recoiling behavior of the cranial tissue. When E11 or E12 cranial tissues (a combination of meningeal/mesenchymal and epidermal sheets), freshly isolated from the otherwise tightly adhesive cerebral hemispheric wall based on differential (shrinking vs. buckling) responses to hypoxia (Figures S3D and S3E), were directly pulled and released, they recoiled, shrinking the cup-like space. See also Figure S4F.

**Figure S4.**
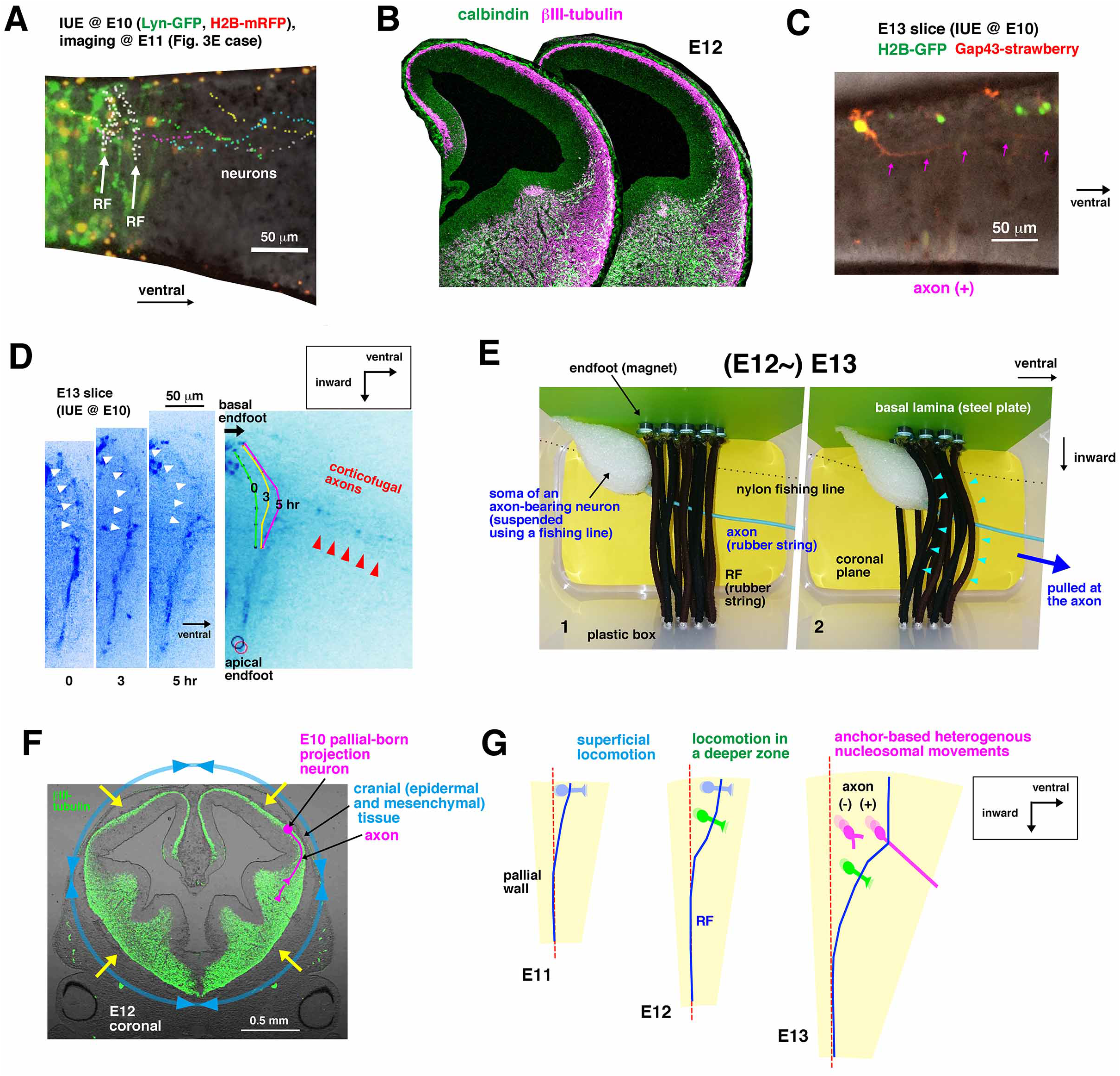
Mechanical factors involved in pushing of RFs by the soma of PP neurons, Related to Figures 3 and 4. (A) Co-tracking of ventrally migrating PP neurons (Figure 3E) and transient ventral shift of two RFs in a coronal slice. (B) Anti–βIII-tubulin and anti-calbindin immunostaining of an E12 hemisphere, showing that calbindin^+^ interneurons originating from the GE use only the most superficial and deepest zones, with minimal (if any) mechanical competition/collision with dorsal-to-ventral migration of E10 pallial-born neurons (Figures S2A and S2B). (C) A PP neuron bearing a ventrally running axon (arrow) in an E13 coronal slice. (D) An E13 coronal slice in which axon-like fibers (red arrowhead) grew ventrally and inward when an RF (white arrowhead), including its basal endfoot, shifted ventrally (Video S6A). (E) Physical model showing that an axon-bearing neuron would push a little deeper part of RFs via its soma. (F) Anti–βIII-tubulin photomicrograph illustrating centripetal constriction by the *in vivo* cranial connective tissue that normally prevents the pallial wall from freely recoiling, reflecting its internal (tissue-intrinsic) mechanical properties that would cause apical curling and basal warping when released from the *in vivo* situation during or after dissection and slicing (Figure 4G). (G) Schematic illustration suggesting that pushing of RFs by the soma of PP neurons occurs in a stage-dependent manner with transitions in the morphology and migratory route of neurons and the depth of RF obliqueness.

**Figure S5.**
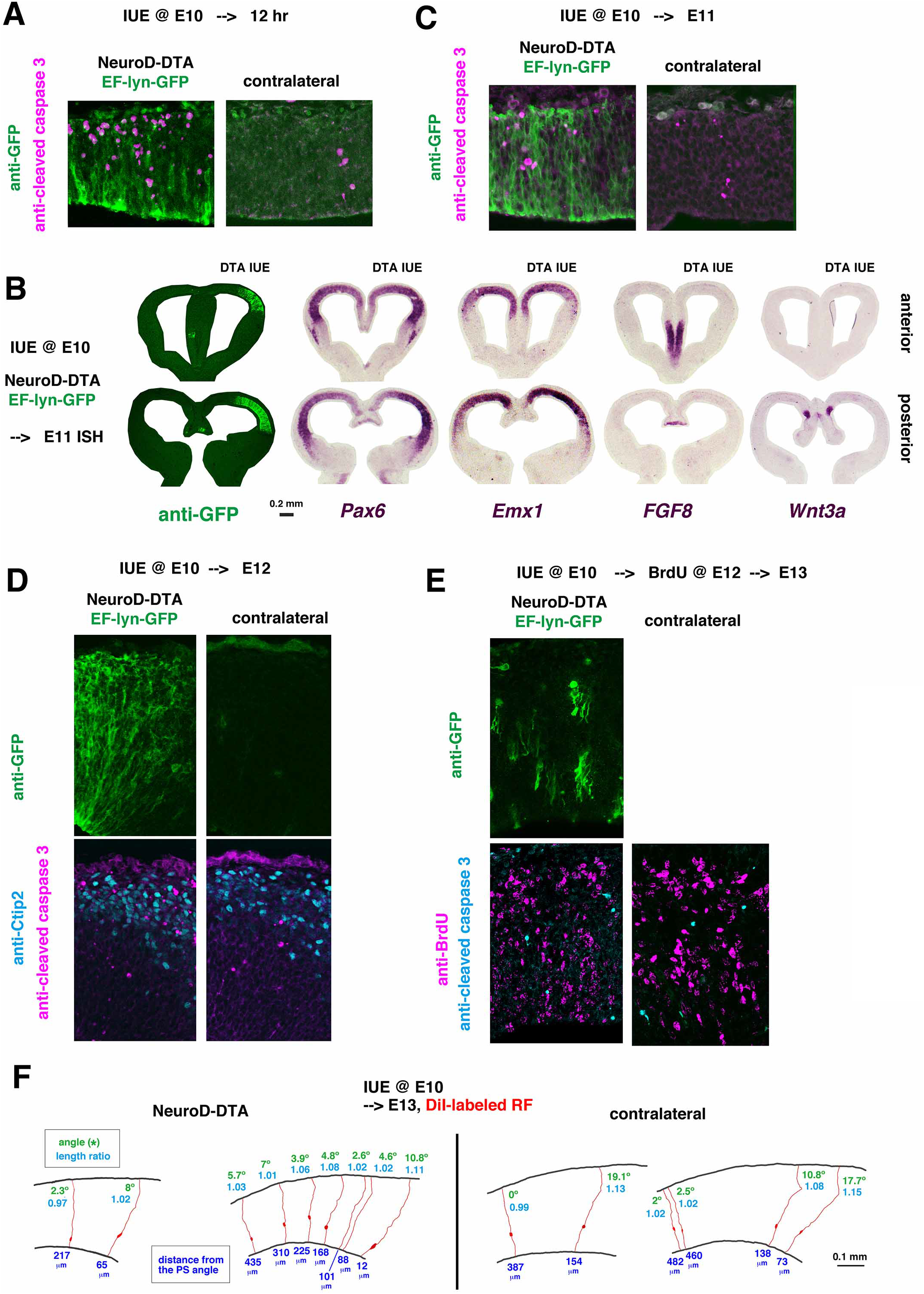
Ventral RF-deflection was reduced by PP depletion via NeuroD-DTA IUE, Related to Figure 5. (A) Anti–activated caspase 3 immunostaining at 12 hr after IUE with NeuroD-DTA and EF-Lyn-EGFP at E10. (B) *In situ* hybridization for Wnt3a, FGF8, Pax6, and Emx1 at E11 following NeuroD-DTA and EF-Lyn-GFP IUE at E10. (C) Anti–activated caspase 3 immunostaining at E11, following IUE with NeuroD-DTA and EF-Lyn-EGFP at E10. (D) Anti–activated caspase 3 and anti-Ctip2 immunostaining at E12, following IUE with NeuroD-DTA and EF-Lyn-EGFP at E10. (E) Anti–activated caspase 3 and anti-BrdU immunostaining at E13, following IUE with NeuroD-DTA and EF-Lyn-EGFP at E10 and BrdU administration at E12. (F) Assessment of RGCs DiI-labeled at E13 in normal (non-IUE) and NeuroD-DTA–IUE hemispheres, following IUE with NeuroD-DTA and EF-Lyn-EGFP. The length ratio and angle of RF deflection are shown. Right, ventral; left, dorsal.

**Figure S6.**
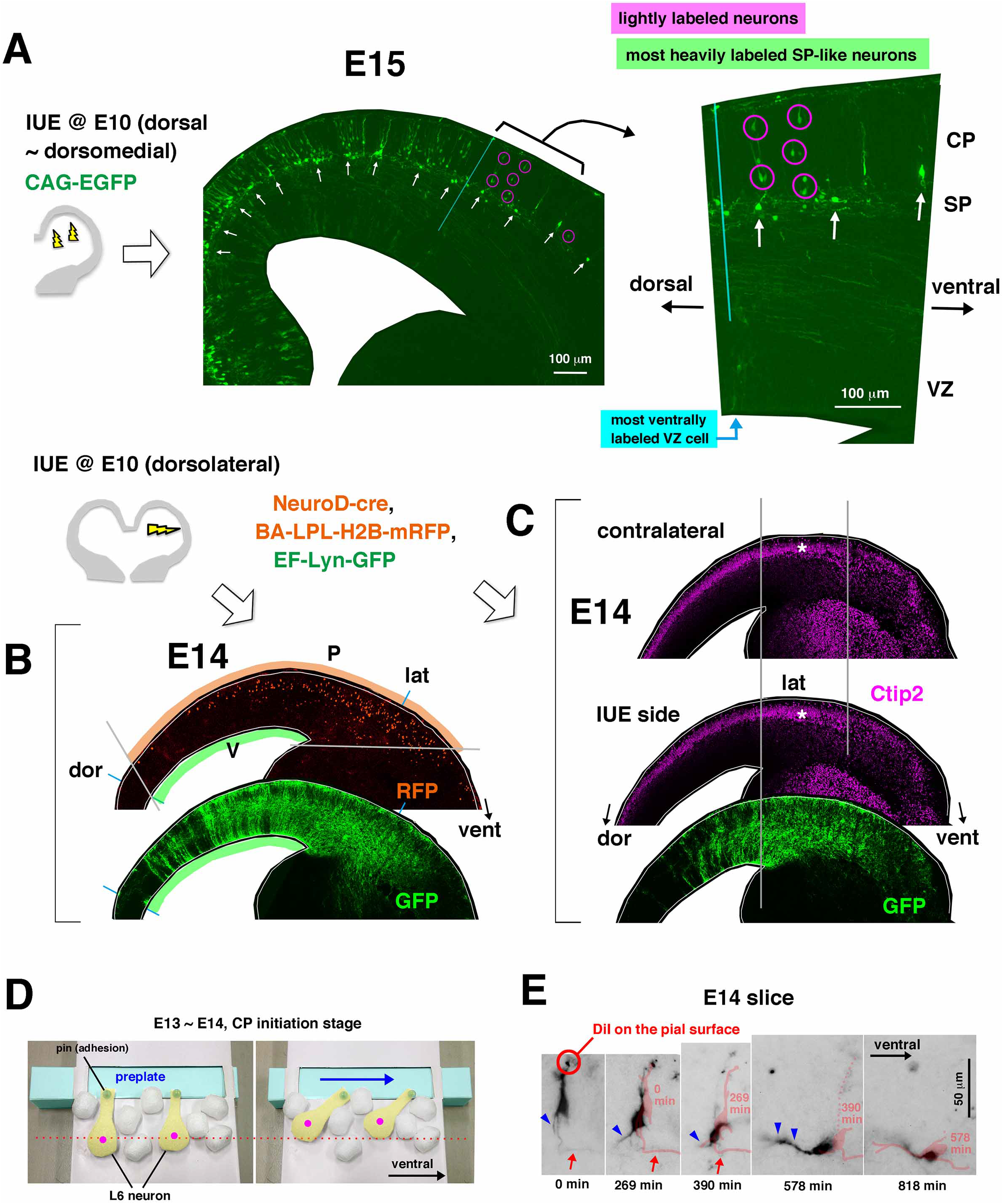
Neuronal behaviors participating in the initiation and the growth of CP, Related to Figure 6. (A) Anti-GFP staining of an E15 hemisphere that had been electroporated (dorsally) at E10. Arrow, most heavily labeled SP-like neurons. Circled, lightly labeled non-SP (layer 6–like) neurons. (B) Anti-RFP and anti-GFP photomicrographs of an E14 cerebral wall after IUE with NeuroD-cre, BA-LPL-mRFP, and EF-lyn-GFP at E10, showing a more ventral distribution of RFP^+^ E10-born neurons in the outer/superficial zone than in the inner GFP^+^ VZ. (C) Anti-Ctip2 (magenta) and anti-GFP (green) photomicrographs of an E14 cerebral wall after IUE with NeuroD-cre, BA-LPL-mRFP, and EF-lyn-GFP at E10, showing that formation of Ctip2^+^ CP was indistinguishable between the electroporated side and the contralateral (un-electroporated) side. (D) Physical model of oblique entrance of neurons destined to contribute to layer 6 into the ventrally streaming PP, by analogy with a person stepping perpendicularly (like L6 neurons) into a horizontally moving walkway (PP-like), which would drag their foot in the direction the walkway is moving. (E) A time-lapse monitored SP-like neuron in an E14 slice. The neuron was labeled with DiI from the pial/outer surface of the wall, and then it retracted its pial process to assume a horizontally extended shape.

**Figure S7.**
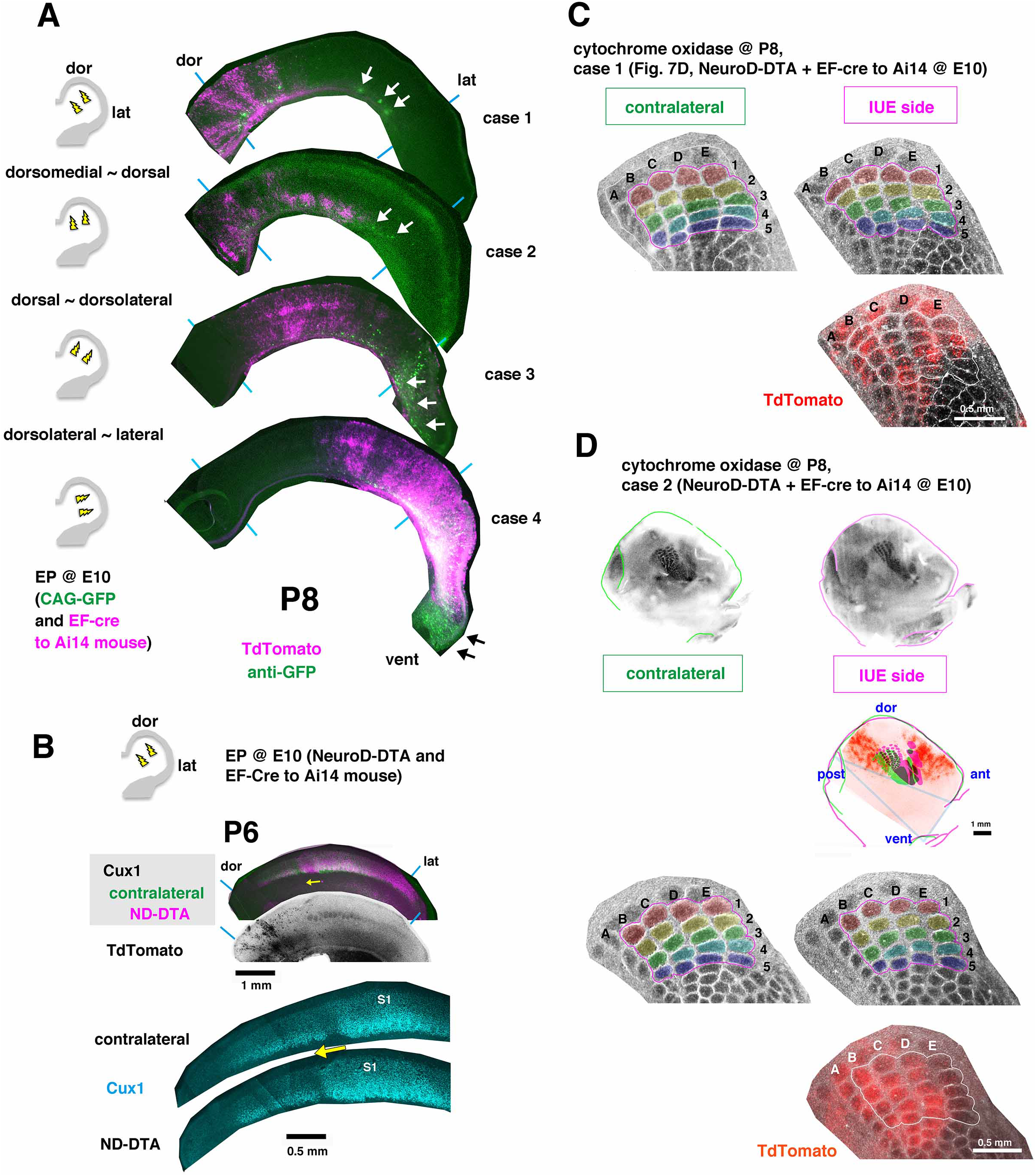
Effects of early embryonic weakening of the PP stream on the postnatal neocortex, Related to Figure 7. (A) TdTomato and anti-GFP photomicrographs of P8 cortices after IUE of Ai14 mice with EF-cre and CAG-GFP vectors. GFP^+^ neurons (arrow) were found more ventrally than in the region most heavily TdTomato^+^. dor, dorsal; vent, ventral; lat, lateral. (B) Anti-Cux1 immunostaining at P6, following IUE of Ai14 mice with EF-cre and NeuroD-DTA vectors at E10. dor, dorsal; lat, lateral. (C, D) High-power comparison of somatosensory barrels of P8 hemispheres between the Neuro-DTA–IUE side and the contralateral side. dor, dorsal; vent, ventral; ant, anterior; post, posterior.

## Legends for Supplemental Movies

### Movie 1, Related to Figure 2

(A) Basal *en face* observation (15 hr) of a cerebral wall flat-mount prepared at E11, following IUE at E10 with pBA-LPL-H2B-mRFP and pEF-cre for 15 hr (summarized in Figures 2A and 2B). Trajectories of neurons that migrated ventrally are superimposed.

(B) Monitoring (8 hr) of a coronally sliced E12 cerebral wall, following IUE at E10 with pBA-LPL-H2B-mRFP and pEF-cre (summarized in Figure 2C).

(C) High-resolution time-lapse monitoring of E10-born PP neurons in an E12 coronal slice, following IUE at E10 with pBA-LPL-H2B-mRFP, pEF-LPL-lynN-EGFP and pEF-cre (summarized in Figure 2D).

### Movie 2, Related to Figure 2

(A) Monitoring (10 hr) of a coronally sliced E13 cerebral wall, following IUE at E10 with pBA-LPL-H2B-mRFP and pEF-cre (summarized in Figure 2E).

(B) Monitoring (15 hr) of a coronally sliced E14 cerebral wall, following IUE at E10 with CAG-EGFP-3NLS (summarized in Figure 2F).

### Movie 3, Related to Figure 3

(A) Time-lapse monitoring of an E12 explant prepared from the PP region and embedded in soft collagen gel containing microbeads (summarized in Figure 3D).

(B) Monitoring (13.6 hr) of an explant prepared from an E11 pallium that had received IUE at E10 (with EF-LPL-Lyn-mCherryy and EF-cre) (summarized in Figure S3A).

(C) Monitoring (240 min) of an E11 coronally sliced cerebral wall cultured in soft collagen gel containing fluorescent microbeads, following IUE at E10

(pBA-LPL-H2B-mRFP, pEF-LPL-lynN-EGFP and pEF-cre) (summarized in Figures 3E and S4A).

### Movie 4, Related to Figure 3

(A) Confocal *en face* observation of an FM4-64-staind E12 pallial wall from the outer surface (summarized in Figure 3F).

(B) High-resolution live monitoring (72.5 min) of three RFs in an E11 pallial wall, following IUE at E10 (with EF-LPL-lyn-EGFP and EF-cre) (summarized in Figure 3G).

(C) Simultaneous monitoring of a single RF (GFP-labeled via IUE at E10) and nuclei of PP neurons in a pallial wall prepared from an E11 H2B-mCherry transgenic mouse (summarized in Figure 3H)

(D) *En face* monitoring (5 hr) of the outer/basal surface of an E11 pallium that had electroporated at E10 (with pBA-LPL-H2B-mRFP, pEF-LPL-lynN-EGFP and pEF-cre) (summarized in Figure S3B).

### Movie 5, Related to Figure 4

(A) Monitoring (5 hr) of a single E10-born neuron in an E13 coronal slice, with tracking of non-axon processes (small arrow) and the nucleus (summarized in Figure 4B).

(B) Live observation (220 min) of a PP neuron retrogradely labeled with DiI inserted in to the internal capsule of an E13 coronal slice (summarized in Figure 4D).

(C) Physical model to simulate a possible pulling force by axons extended corticofugally to the subpallium (summarized in Figure 4F). Elastic property given to the axon using a rubber string causes a warping response of the pallial wall.

(D) Outward warping response shown by a freshly prepared E13 coronal cerebral slice (by 119 min), following the initial apical curling of the pallial part occurred by 19 min. See also Figure 4G.

### Movie 6, Related to Figure 6

(A) Monitoring (5 hr) of an E13 coronally sliced cerebral wall, following IUE at E10 with BA-LPL-Gap43-strawberry and EF-cre (summarized in Figure S4D). When axon-like fibers (red arrowhead) grew ventrally and inward, a RF (white arrowhead), including its basal endfoot, shifted ventrally.

(B) A coronal slice prepared from an E14 H2B-mCherry mouse, monitored for 6.9 hr (summarized in Figure 6A).

## Acknowledgement

We thank T. Shimogori for instructing *in utero* electroporation at E10 and comments to the manuscript; T. Kawauchi, R.L. Huganir, F. Matsuzaki, K. Kaibuchi, T. Sapir for plasmids; M. Ogawa for suggestions; M. Masaoka, T. Sato, and Y. Muto for excellent technical assistance; Miyata lab’s people for discussion. This work was supported by MEXT Grant-in-Aid for Scientific Research on Innovative Areas “Cross-talk between moving cells and microenvironment as a basis of emerging order” 22111006 (T.M.), JSPS Grant-in-Aid for Scientific Research (A) 16H02457 (T.M.), and JSPS Grant-in-Aid (C) 17K10176 (K.S.).

## Author Contributions

K.S. and T.M. designed the study, and acquired and analyzed the data. K.S., M.O., N.N. performed the overall experiments, with DTA-based experiments by Y.W., postnatal analyses by Y.N., mechanical assays by A.N., and live imaging by T.S. and A.S.. T.M. wrote the manuscript, with input from the other authors.

## Methods

### Animals

Animal experiments were conducted according to the Japanese Act on Welfare and Management of Animals, Guidelines for Proper Conduct of Animal Experiments (published by Science Council of Japan), and the Fundamental Guidelines for Proper Conduct of Animal Experiments and Related Activities in Academic Research Institutions (published by the Ministry of Education, Culture, Sports, Science and Technology of Japan). All protocols for animal experiments were approved by the Animal Care and Use Committee of Nagoya University (No. 29006). R26-H2B-mCherry transgenic mice (accession No. CDB0239K) (Abe et al., 2011) were provided by Toshihiko Fujimori (NIBB, Japan). Ai14 mice were purchased from Jackson Laboratory (007914). Timed-pregnant ICR mice were obtained from SLC (Hamamatsu, Japan). *Reeler* mice (gift from Dr. Masaharu Ogawa, RIKEN) and CXCR4 knock-out mice (Tachibana et al., 1998) (gift form Dr. Takashi Nagasawa, Osaka University) were maintained and used in Nagoya University. The day when the vaginal plug was detected was considered E0.

### *In utero* electroporation

*In utero* electroporation (IUE) of E10 mouse embryos was performed as described previously (Okamoto et al. 2013; Otaka-Maruyama et al., 2018). After DNA solution was injected into the lateral ventricle, the head of the embryo in the uterus was placed between the discs of a forceps-type electrode (disc electrodes of 1 mm; CUY560P1, NEPA GENE, Chiba, Japan), and electric pulses (50 V) were administered four times, resulting in gene transfection into the cerebral wall.

### Plasmids

For visualization of preplate neurons, E10 mice were electroporated with a mixture of pEF-LPL-Lyn-EGFP (0.5 µg/µl) and/or pBA-LPL-H2B-mRFP1 (0.5 µg/µl) and a Cre-recombinase expression plasmid, pEF-Cre (0.001 µg/µl for sporadic labeling; 0.1 µg/µl for massive/diffuse labeling) (Okamoto et al., 2013; Sakakibara et al., 2014) (Figures 2A, 2B, 2C, 2D, 2E, 3E, 3G, 3H, S3A, S4A) or pNeuroD-Cre (Sapir et al., 2008) (Figures S2A, S2B, S2C, S2G, S2I, S6B). Alternatively, a mixture of pBA-LPL-GAP43N-mStrawberry (0.5 μg/μl) (constructed from pH2B-Strawberry [Addgene], Gap43 N-terminal membrane-targeted signal sequence, and pBA-LPL-H2B-mRFP) and pCAG-EGFP-3NLS (0.5 µg/µl) (Konno et al., 2008) (Figures 2F, 4B, S2E, S2F, S4C) was used. In other experiments, pEF-LPL-Lyn-mCherry (Watanabe et al., 2018) (Figure S4A), pCAG-EGFP (0.5 μg/μl) (Figures 7B, S2D, S2H, S6A, S7A), or pEF-Lyn-EGFP (0.5 μg/μl) was used (Figures 4G, 4H, 5B, 6C-6K, S2C, S5A–S5E, S6B, S6C). For IUE-mediated neuron ablation, E10 ICR mice were electroporated with pNeuroD-DTA (1 μg/μl) together with pEF-LPL-Lyn-EGFP (0.5 µg/µl) and pEF-Cre (0.1 µg/µl), as previously described (Watanabe et al., 2018) (Figures 5B–E, 6C–K, S5A–S5E). pNeuroD-DTA IUE was also performed on Ai14 mice with pEF-Cre (0.1 µg/µl) (Figures 7C, 7D, S7C, S7D). pEF-LPL-N17-Rac1 (1 μg/μl), constructed from pEF-BOS-N17-Rac1 (Kuroda et al., 1998) and pEF-LPL-Lyn-EGFP, and pEF-LPL-DN-N-Cadherin (1 μg/μl), constructed from pTα1-DN-N-Cadherin (Kawauchi et al., 2010; Nuriya and Huganir et al., 2006) and pEF-LPL-Lyn-EGFP, were used with NeuroD-Cre (1 μg/μl) (Figure S2I).

### DiI labeling

For retrograde labeling of PP neurons, DMEM/F12 containing a high concentration of extremely fine crystals of 1,1-dioctadecyl-3,3,3,3-tetramethylindocarbocyanine perchlorate, DiI C18(3) (D-282; Molecular Probes) was injected into coronal slices of E13 hemispheres (Figures 4C and 4D). For DiI labeling from the pial surface, pallial walls were incubated for 3 min in a highly diluted suspension, which was made by adding 5–10 μl of DiI stock solution (10 mg/ml in ethanol) to 10 ml DMEM/F12, as described previously (Miyata et al., 2001) (Figure S6B).

### Live imaging

Cross-sectional or *en face* cultures of cerebral walls were prepared manually using a microknife and a silicone rubber plate, as described previously (Miyata et al., 2001; Okamoto et al., 2013; Shinoda et al., 2018). Briefly, cerebral walls were microsurgically processed and embedded in a polystyrene cell-culture dish (Corning) coated with type-I collagen gel (Nitta Gelatin) at a concentration of 0.9–1.1 mg/ml, and cultured in DMEM/F12 supplemented with 5% horse serum, 5% fetal bovine serum, N2 (1:100, Thermo Fisher Scientific), EGF (10 ng/ml, Peprotech), and bFGF (10 ng/ml, Gibco).

Time-lapse confocal microscopy was performed on an inverted CV1000 spinning disc confocal system (Yokogawa, Japan) with a 40× objective lens (UPLFLN 40XD, (NA) 0.75, Olympus, Japan), a BX51W1 microscope (Olympus) equipped with a CSU-X1 spinning disc confocal unit (Yokogawa) with a 40× objective lens (LUMPLFLN40XW, Olympus) or 100× objective lens (LUMPLFL100XW, Olympus) and an iXon+ EMCCD camera (Andor), in an on-stage culture chamber (Tokai Hit) filled with 55% N_2_, 40% O_2_, and 5% CO_2_. For confocal imaging of all cells near the pial surface (Figure 3F), cerebral hemispheric walls were stained with FM4-64 (Thermo Fisher Scientific) at a concentration of 5 μg/ml, as described previously (Shinoda et al., 2018). Monitoring of explants on culture dishes (Figure S3A) was performed using an inverted CV1000 system (Yokogawa, Japan) with a 20× objective lens.

### Assay on cranial tissues

For hypoxic treatment of embryos at E10–12, excised uteri containing embryos were placed in 60-mm culture dishes containing culture medium and placed in a CO_2_ incubator (37°C, 20% O_2_). After 1–2 hr, the embryos were fixed and sectioned, showing physical separation between the cranial soft tissue (composed of the meningeal mesenchyme and the overlying epidermis) and the otherwise tightly adhesive cerebral wall (Figures S3D and S3E). Live cranial tissues isolated through such hypoxia-inducing incubation were then transferred into oxygenized culture medium and further subjected to direct mechanical assessment under a dissection microscope (Figure S3F).

### Slice bending assay

Coronal slices (250–300 μm thick) were manually prepared from E13 cerebral hemispheres (both pallial and striatal regions together), and then placed into a polystyrene suspension cell culture dish (Corning). Bending and/or warping (Figures 4G, 4H) was recorded using an Olympus IX71 (4×) equipped with an on-stage culture chamber (Tokai Hit), temperature-controlled and filled with 5% CO_2_, and an Orca ER camera (Hamamatsu Photonics), as previously described (Okamoto et al., 2013; Shinoda et al., 2018). In some slices, transections were made at the route of corticofugal axons (Figure 4H).

### Beads traction assay

Explants (Figure 3D) or slices (Figure 2E) were embedded very soft (0.2–0.3 mg/ml) collagen gel containing red fluorescent microbeads (Fluoro-Max [Thermo Fisher Scientific], 1 μm in diameter, 0.5% v/v).

### Physical simulations

Based on the usefulness of physical (non-mathematical) models for simulating morphogenetic events during development (Davis, 2013), simulations of physical events involving PP neurons (Figures 4F, S3C, S4E, S6D) were prepared using mechanically appropriate materials (e.g., rubber strings to mimic RFs [Shinoda et al., 2018] and axons [Bray,1984; Lamoureux et al., 1989] and styrofoam blocks to model neurons).

### Immunohistochemistry

Cross-sectional and whole-mount immunohistochemistry experiments were performed as described previously (Okamoto et al., 2013; Shinoda et al., 2018). Embryonic brains were fixed for 1 hr at 4°C with periodate–lysine–paraformaldehyde (PLP) fixative. For immunostaining of phospho-myosin light chain, 2% trichloroacetic acid in PLP was used. Frozen sections (16 μm thick) were treated with the following primary antibodies: 9-4c (mouse, gift from Tatsumi Hirata, hybridoma supernatant), anti-Pax6 (rabbit, PRB-278O, COVANCE, 1:300), anti-Tbr1 (rabbit, ab31940, Abcam, 1/500), anti-Nestin (rabbit, gift from Yasuhiro Tomooka, 1:2000), RC2 (mouse IgM, gift from Miyuki Yamamoto, hybridoma supernatant), anti–Laminin A chain (rabbit, L9393, Sigma-Aldrich, 1:50), anti-L1 (rat, MAB5272, Millipore, 1:200), anti-MAP2 (mouse, M1406, Sigma-Aldrich, 1:5000), anti–cleaved caspase 3 (rabbit, #9661, Cell Signaling, 1:200), anti–phospho-myosin light chain (ab2480, Abcam, 1:500), anti-calbindin (mouse, C9848, Sigma, 1:3000), anti-TAG1 (goat, AF1714, R&D Systems, 1:400), anti–βIII-tubulin (mouse, MMS-435P, COVANCE, 1:1000), anti-BrdU (rat, NB500-169, Novus Biologicals, 1:2000) (following intraperitoneal administration of BrdU [50 μg/g b.w.] to a pregnant female mouse), anti-Ctip2 (rat, ab18465, Abcam, 1:1000); anti-Satb2 (mouse, ab51502, Abcam, 1:400); anti-Cux1 (rabbit, sc-13024, Santa Cruz Biotechnology, 1:300), anti-GFP (rat, 04404-84, Nacalai Tesque, 1:500; rabbit, 598, MBL, 1:500; chick, GFP-1020, Aves Labs, 1:500). After washes, sections were treated with Alexa Fluor 488–, Alexa Fluor 546–, or Alexa Fluor 647–conjugated secondary antibodies (Molecular Probes, A-11029, A-11006, A-11034, A-11030, A-11035, A-11081, A-21236, A-21245, 1:200), and subjected to confocal microscopy (FV1000; Olympus, Tokyo, Japan).

### *In situ* hybridization

Riboprobes were synthesized using the following primers: Emx1 (703-1118, NM_010131) Fw: TACAAACGGCAGAAGCTGGAA, Rv: TATTCCCATAGGGAAGGGGGA; Pax6 (702-1547, NM_001244198) Fw: AGGGGGTCTGTACCAACGAT, Rv: ACATGTCAGGTTCACTCCCG; Wnt3a (1805-2643, NM_009522) Fw: TGAAGACTCATGGGATGGAGC, Rv: GTCTAAATCCAGTGGTGGGTGG; FGF8 (537-1009, XM_006526668) Fw: ATCCGGACCTACCAGCTCTAC, Rv: TATGCACAACTAGAAGGCAGCTC; Rorβ (1107-1994, NM_001043354) Fw, GACATGACTGGGATCAAACAGAT; Rv, GCCTTGTACAATGGAGGAAACAGT. Probes were labeled with digoxigenin (Roche Diagnostics, Sigma-Aldrch). E11 cerebral hemispheres that were electroporated at E10 with NeuroD-DTA were fixed overnight with 4% paraformaldehyde at 4°C. Postnatal mice that had been electroporated at E10 with NeuroD-DTA were fixed by transcardiac perfusion with 4% paraformaldehyde. Frozen sections (20 μm) were pretreated with 4% paraformaldehyde (5 min) and 0.1 M triethanolamine / 0.25% acetic anhydride (10 min), and then hybridized at 60°C for 15 hr in a solution containing 50% formamide, 0.1% SDS, 0.64 mg/ml tRNA, and 0.5 μg/ml riboprobes. The labeled cRNAs were visualized using nitroblue tetrazolium/5-bromo-4-chloro-3-indolyl phosphate solution.

### Scanning electron microscopy

Cerebral hemispheres were fixed with 4% paraformaldehyde / 2.5% glutaraldehyde in 0.1 M phosphate buffer (PB) at 4°C, trimmed with a blade, and then subjected to post-fixation in 1% osmium tetroxide in 0.1 M PB. The samples were dehydrated and coated with osmium. Images were collected on a JSM-7610F Schottky Field Emission SEM (JEOL).

### Quantification and statistical analysis

Image analyses and quantifications were conducted using ImageJ (National Institute of Health). Nuclear tracking was performed using the Move-tr/2D software (Library Co., Ltd., Japan). The length ratio for each DiI-labeled RGC (Figure 5D) was obtained by dividing the total length of a RGC by the length of a straight line drawn perpendicularly from the apical/ventricular endfoot of that RGC to the outer/pial surface of the wall. The angle (Figure 5E) was measured between the intra-VZ portion of an RGC and a straight line drawn perpendicularly from the apical endfoot of that RGC. To evaluate correlations between the length ratio and the angle with the distance from the PS angle, the Spearman rank correlation coefficient was calculated (Figures 5D and 5E). Differences were analyzed using the Exact Wilcoxon rank–sum test (Figures 4G, 4H, 6G, 6H, 6I) and ANCOVA (Figures 5D and 5E) in the R statistical environment. All values represent means ± standard error of the mean (SEM). Significance was set at p=0.05. No statistical methods were used to predetermine sample size, but sample sizes were similar to those described in related studies (Okamoto et al., 2013; Shinoda et al., 2018). No randomization of samples was performed, and no blinding was performed. The number of samples or cells examined in each analysis is shown in the legends or text.

**Movies (S1-S6).**
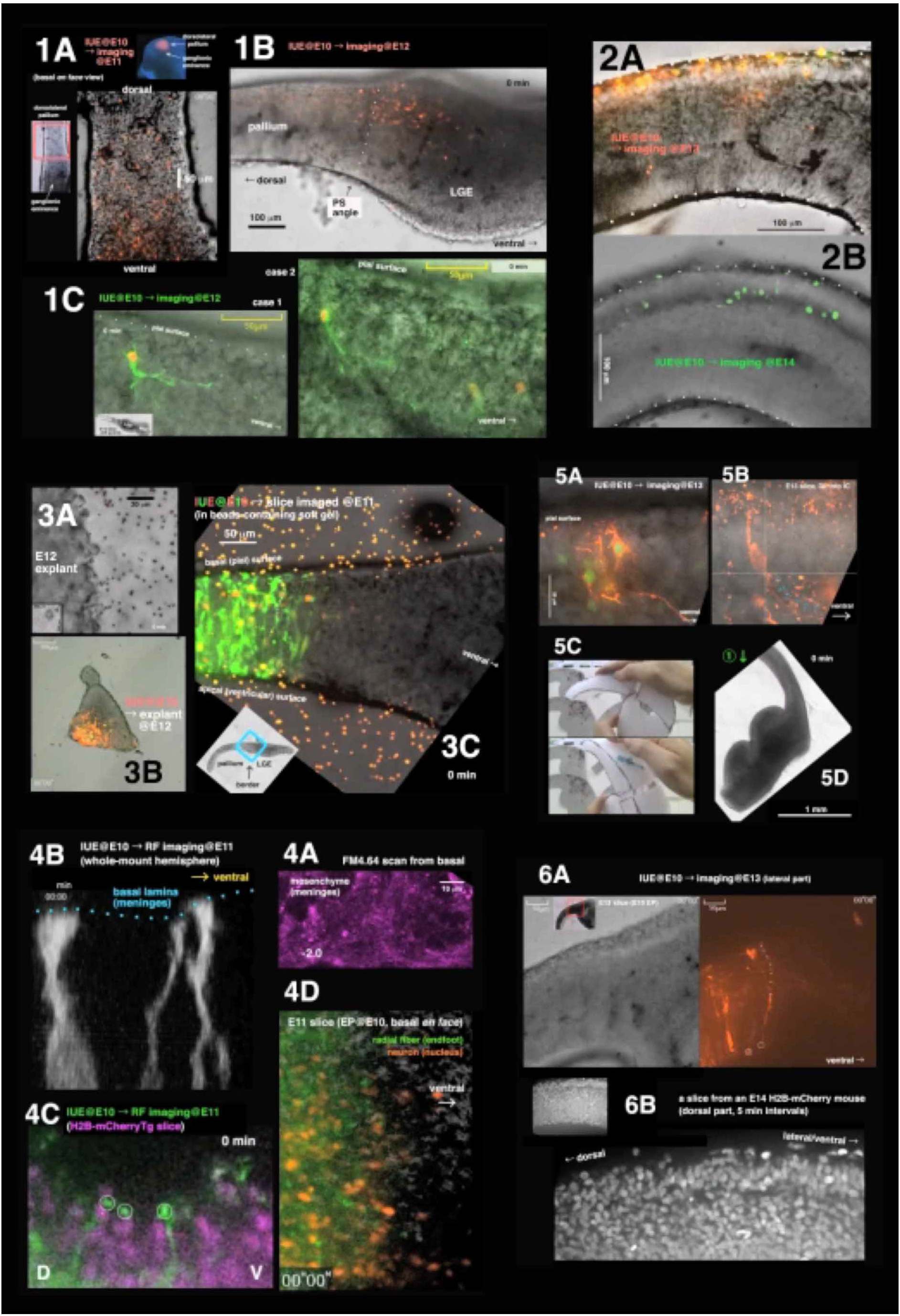

